# Synthetic signaling platform uncovers and rewires cellular responses to PD-1 perturbation

**DOI:** 10.1101/2025.10.01.679874

**Authors:** Zhixing Ma, Lars Hellweg, Susanna K. Elledge, Jasper B. Lee, Maria Caterina Rotiroti, Kai W. Wucherpfennig, Robbie G. Majzner, Xin Zhou

## Abstract

Tyrosine phosphorylation motifs are central regulators of cell signaling, yet methods to selectively detect and reprogram these events have been lacking. Here we introduce Sphyder (Selective PHosphotYrosine DEtection and Rewiring), which enables precise detection of signaling at the resolution of individual phosphorylation motifs. Using Sphyder biosensors, we resolved phosphorylation dynamics and uncovered regulatory mechanisms of the checkpoint receptor PD-1 in living cells. Sphyder also provided a framework for reconstructing phosphosignaling pathways. With this approach, we redirected PD-1 signaling from immunosuppressive to immunoactivating outputs and engineered synthetic receptors that linked extracellular sensing to customized transcriptional programs. In addition, Sphyder biosensors revealed previously unrecognized mechanisms of the PD-1/VEGF bispecific antibody Ivonescimab, showing that it induces VEGF-dependent clustering, phosphorylation, and degradation of PD-1. These findings may underlie its promising clinical activity relative to conventional PD-1 blockade. Together, our study establishes a broadly applicable strategy for sensing and reprogramming cell signaling, while also providing mechanistic insights into a new class of immune checkpoint inhibitors of major clinical interest.

## Introduction

Tyrosine phosphorylation is a central mechanism by which cells sense and respond to extracellular cues (*1*). In T cells, signaling motifs containing tyrosine residues act as modular switches that recruit downstream effectors when phosphorylated (*2*). The output of cell signaling is therefore determined by which motifs are engaged and their phosphorylation state. For example, the programmed cell death receptor PD-1 contains an immunoreceptor tyrosine-based switch motif (ITSM) that, once phosphorylated, recruits the phosphatase SHP2 and suppresses T cell activation (*3*, *4*). By contrast, phosphorylation of the adaptor protein LAT recruits signaling proteins that propagate T cell receptor signaling (*5*). Together, such phosphorylation events create a modular network that enables precise regulation of cellular responses.

Significant progress has been made in mapping tyrosine phosphorylation motifs and identifying their downstream effectors. Yet tools that can probe or manipulate phosphorylation at the level of individual motifs in living cells remain scarce, limiting our ability to dissect their spatiotemporal regulation. Canonical phosphotyrosine-binding domains such as Src-homology 2 (SH2) domains typically exhibit modest affinity and rely on avidity-based interactions rather than strict sequence selectivity (*6–9*). Engineered SH2 “superbinders” improve affinity but remain promiscuous in binding (*10*, *11*). Fluorescent kinase sensors report on global kinase activity but do not resolve the phosphorylation state of specific motifs (*12*), while phospho-specific antibodies provide higher selectivity but are generally incompatible with intracellular applications (*13–15*). Thus, despite extensive biochemical knowledge, new approaches for site-resolved analysis of signaling in living cells are needed (*16*). Moreover, strategies that allow site-specific perturbation and rewiring would provide powerful tools to both interrogate and reprogram immune signaling.

Phosphotyrosine signaling is especially critical for programmed cell death protein 1 (PD-1), an immune checkpoint receptor of significant clinical relevance (*17*). Antagonist antibodies such as Pembrolizumab and Nivolumab have transformed cancer immunotherapy, while agonist antibodies are being explored in autoimmune and inflammatory diseases (*18*). Despite these advances, PD-1 signaling is not fully understood. For instance, PD-1 was long considered a monomeric receptor, but recent work suggests that dimerization contributes to its activity (*19*), though how this influences PD-1 phosphorylation and receptor function remains unclear. At the same time, therapeutic strategies that move beyond conventional PD-1 blockade are emerging. The PD-1/vascular endothelial growth factor (VEGF) bispecific antibody Ivonescimab has shown encouraging outcomes in clinical studies (*20*). However, the molecular basis for this distinction remains undefined. In particular, it is unclear whether Ivonescimab affects PD-1 signaling in a different manner than other PD-1/PD-L1 inhibitors, or whether its activity exceeds that of combined PD-1 and VEGF pathway inhibition (17). Addressing such questions requires tools capable of resolving receptor functional states in living cells.

Here, we report Sphyder (Selective PHosphotYrosine DEtection and Rewiring), a platform for engineering nanomolar-affinity binders with strict sequence selectivity for individual phosphosites. Unlike native recognition modules, Sphyder modules function as orthogonal, site-specific signaling units. This capability enables two complementary applications: biosensors that allow real-time visualization of phosphorylation dynamics, and synthetic circuits that rewire phosphorylation inputs into new functional outputs. Utilizing PD-1 as a model, Sphyder biosensors revealed previously undercharacterized features of receptor regulation, including the contribution of dimerization to PD-1 activation. Crucially, they provided the first live-cell view of the PD-1/VEGF bispecific antibody Ivonescimab, revealing that it induces VEGF-dependent clustering, phosphorylation, and degradation of PD-1, a mechanism distinct from conventional PD-1 blockade.

Beyond sensing, Sphyder circuits redirected PD-1 phosphorylation from inhibitory signaling through SHP2 recruitment to activating pathways that drive kinase activation and cytokine expression, as well as enabled synthetic receptor engineering. When incorporated into Chimeric Antigen Receptor (CAR)-T cells, these rewired signaling modules enhanced anti-tumor activity. Importantly, the modular design was not restricted to PD-1. Extension of Sphyder to LAT (Linker for Activation of T cells), an adaptor protein that facilitates the transmission of signals from the TCR to downstream signaling pathways, demonstrated its broader applicability across cell signaling pathways. Collectively, these findings highlight Sphyder as a versatile approach for uncovering regulatory mechanisms of cell signaling—including the first live-cell mechanistic elucidation of Ivonescimab—and for building new strategies in therapeutic cell engineering.

## Results

### Engineering a site-specific protein binder to PD-1 pY248

To engineer a high-affinity and sequence-specific binder for PD-1 pY248, we employed an intramolecular phosphopeptide sandwich strategy. The C-terminal SH2 domain (C-SH2) of the SHP2 phosphatase binds the phosphorylated ITSM motif (Y248) of PD-1 with a dissociation constant (Kd) of ∼10 µM (*6*) (*21*). This low-affinity interaction initiates the recruitment of SHP2, which is then followed by binding of the N-terminal SH2 domain (N-SH2) to PD-1 pY223 (*22*). These interactions drive SHP2 activation, resulting in dephosphorylation of immune receptors such as CD28 and suppression of T cell activity (*4*). We hypothesized that the SHP2 C-SH2– PD-1 pY248 complex could present a composite epitope for nanobody (Nb) recognition. By linking an Nb to the SH2 domain, we could generate a binder with high affinity and sequence selectivity, in which phosphospecificity is conferred by the SH2 domain and affinity and sequence specificity are mediated by the Nb (**Fig. 1A**)

**Figure 1.**
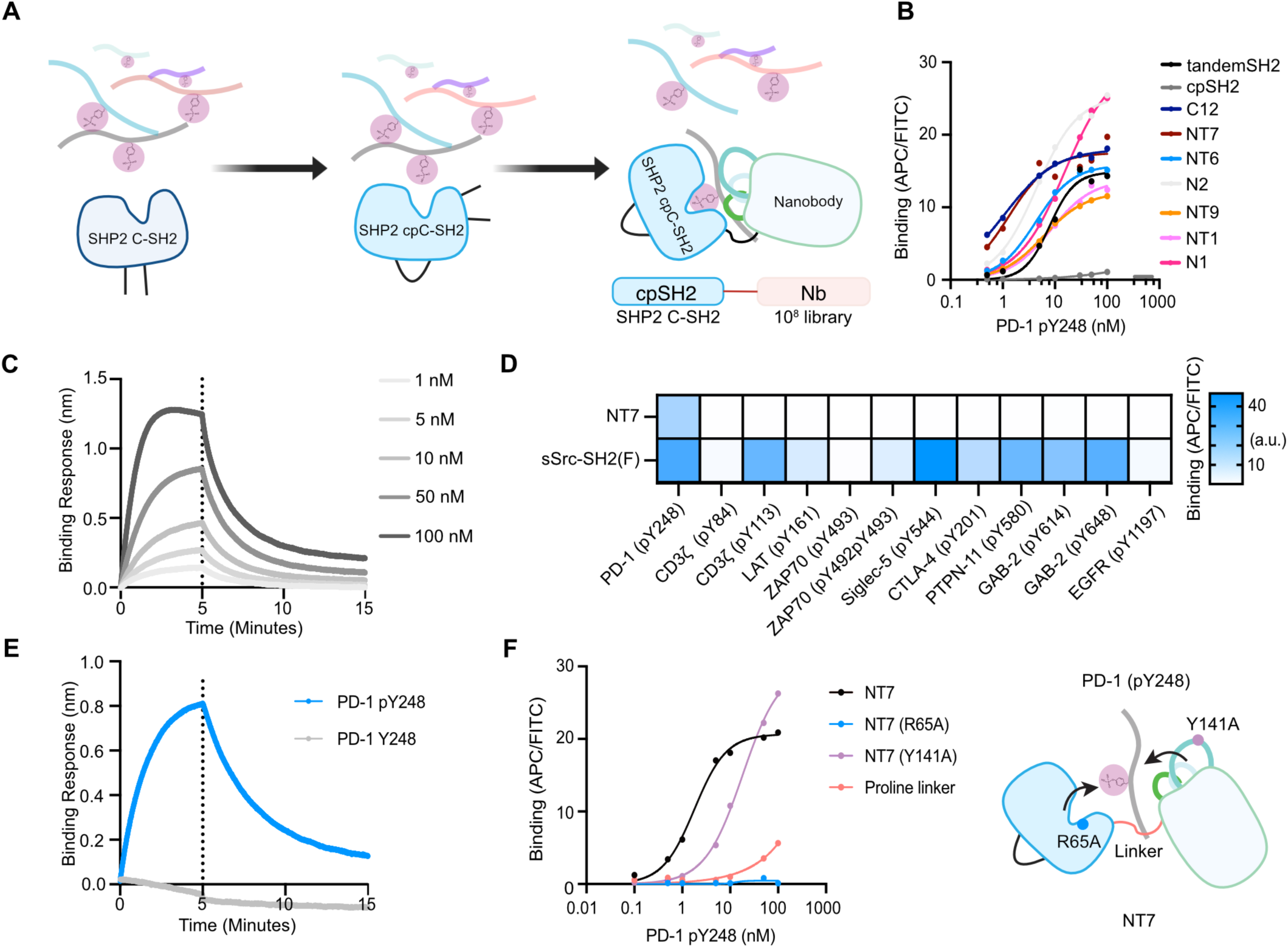
PD-1 pY248 binder engineering and biochemical characterization. **(A)** Schematic of the engineering strategy for generating high-affinity, site-specific binders against phosphorylated PD-1. A yeast surface display library was constructed by fusing the SHP2 C-terminal SH2 domain to a Nb library to create a two-domain architecture for high-affinity and high-specificity phosphopeptide binding. **(B)** Flow cytometry analysis of yeast-displayed PD-1 binders showing binding titrations against biotinylated PD-1 pY248 phosphopeptide. Data are shown for cpSH2 alone, a tandem SH2 domain, and several selected nanobody fusions (C12, NT7, NT6, N2, NT9, NT1, N1). Data represent fluorescence quantification of binding (APC signal) normalized to expression (FITC signal) from 10,000 yeast cells. **(C)** BLI binding analysis of binder NT7 showing concentration-dependent binding curves for PD-1 pY248 peptide at indicated concentrations (1–100 nM). The calculated Kd is 17.2 ± 8.21 nM. **(D)** Heat map showing relative binding strength of the engineered PD-1 phosphosite binder NT7 displayed on yeast against a panel of phosphotyrosine peptides from diverse immune signaling proteins. Data represent mean fluorescence quantification of binding (APC signal) normalized to expression (FITC signal) from n = 2 replicates and normalized to no incubation with peptides. **(E)** BLI binding kinetics demonstrating specificity of 100 nM NT7 binder for PD-1 pY248 versus unphosphorylated PD-1 peptides. **(F)** Flow cytometry titrations comparing binding of the NT7 clone and NT7 mutants or linker variants displayed on yeast to PD-1 pY248 peptide, indicating effects on binding affinity. Right is the representative schematic of NT7 and mutants. Data represent fluorescence quantification of binding (APC signal) normalized to expression (FITC signal) from 10,000 yeast cells.

To test this hypothesis, we first engineered a circularly permuted C-SH2 (cpSH2) domain, in which the native termini were joined, and new termini introduced at alternative positions to allow close spatial proximity of the phosphobinding site to the fused Nb (**Fig. 1A, Fig. S1A**). Of three candidate permutations, the EF-loop breakpoint yielded the most stable cpSH2 domain, though with reduced affinity for PD-1 pY248 compared to wild-type SH2 (**Fig. S1B**). We next constructed a yeast surface display library encoding cpSH2 fused via a short flexible linker (GGSGGS) to a diversified Nb domain (*23*). Library diversity was ∼10^8^, with expected amino acid distributions across complementarity-determining regions (CDRs; **Fig. S2A**). One round of magnetic selection and four rounds of fluorescence-activated cell sorting (FACS) using biotinylated PD-1 pY248 peptide with decreasing peptide concentrations per round enriched for higher-affinity binders (**Fig. S2B**). One round of negative selection against unphosphorylated peptide was performed to ensure phosphospecificity of the identified binders.

After the final selection round, NGS was performed and the seven most-abundant sequences were synthesized and characterized in yeast display assays for binding to PD-1 pY248 peptide. The binders demonstrated nanomolar binding EC_50_ and substantially enhanced binding relative to cpSH2 alone, with clones NT7 and C12 showing the lowest EC_50_s (**Fig. 1B**). NT7 and C12 have approximately 10-fold improved EC_50_s compared to the tandem SH2 construct. Specificity profiling against a panel of immune phosphotyrosine-containing peptides abundant in T cells showed that the binders were highly selective for phosphorylated PD-1 (**Fig. S2C**).

The top-performing binder NT7 was purified as a recombinant protein for further characterization. Biolayer interferometry (BLI) measurements showed that NT7 bound phosphorylated PD-1 with a dissociation constant of 17 nM, approximately 1,000x tighter than binding of the N- or C-SH2 domains in SHP2 (**Fig.1C**). NT7 showed no binding to unphosphorylated PD-1 peptide or phosphopeptides from other immune receptors and proteins, including CD3ζ pY84, CD3ζ pY113, LAT pY161, ZAP-70 pY493, ZAP-70 pY492pY493, Siglec-5 pY544, CTLA-4 pY201, PTPN-11 pY580, GAB-2 pY614, GAB-2 pY648, and EGFR pY1197 (**Fig.1C-E, Fig. S2**). Notably, although Siglec-5 contains an ITSM motif highly similar to PD-1, the selected binders retained exclusive specificity for phosphorylated PD-1 (**Table S1**).

To elucidate the binding mechanism of NT7, we performed mutational analyses. Substitution of R65, which corresponds to the phosphotyrosine recognition site of SH2 domains with alanine completely abolished binding (**Fig. 1F, Fig. S2E**). Substitution of the flexible glycine–serine linker connecting cpSH2 and the Nb with a rigid proline-rich linker markedly reduced binding, indicating that conformational cooperativity between cpSH2 and the Nb contributes to affinity (**Fig. 1F**). In addition, mutation of Y141 in the CDR3 loop of Nb, which AlphaFold3 predicted to contact the peptide, to alanine decreased affinity nearly 10-fold relative to the wild type (**Fig. 1F**). These results demonstrate that NT7 achieves high-affinity and highly specific recognition of PD-1 pY248 through engagement of both the engineered SH2 domain and the Nb.

### Live-cell PD-1 activity sensors reveal effects of receptor dimerization

To assess whether NT7 binds PD-1 pY248 in a phosphorylation-dependent manner in mammalian cells, we generated HEK293 cells co-expressing NT7-mCherry and GFP-tagged PD-1 variants, either wild-type or containing tyrosine-to-phenylalanine mutations. Cells were also engineered to express lymphocyte-specific protein tyrosine kinase (LCK) which phosphorylates PD-1. In cells expressing wild-type PD-1, NT7-mCherry localized to the plasma membrane, indicating binding to phosphorylated PD-1 (**Fig. S3A**). NT7-mCherry also retained membrane localization in cells expressing the Y223F mutant. In contrast, no membrane localization was observed in cells expressing Y248F or Y223F/Y248F mutants. These results confirmed that NT7 binds specifically to phosphorylated Y248 in PD-1.

Fluorescence recovery after photobleaching (FRAP) analysis was performed to assess the dynamics of protein binding in cells. PD-1-CFP exhibited slow fluorescence recovery, indicating limited exchange of receptors once the immune synapse is formed. In contrast, NT7-YFP showed rapid signal recovery, suggesting that NT7 dynamically dissociates and reassociates with phosphorylated PD-1 **(Fig. S3B)**. This feature supports the application of NT7 to report real-time phosphorylation kinetics of PD-1.

To characterize the dynamics of PD-L1-induced PD-1 phosphorylation with spatial information, we developed a confocal imaging–based assay using Jurkat T cells expressing PD-1-CFP and NT7-YFP, and Raji B cells expressing PD-L1-mCherry. Raji cells were first immobilized on the surface of a flow-cell chamber, followed by the introduction of Jurkat cells using a syringe-based flow system. In the presence of the anti-CD3/anti-CD19 bispecific T cell engager (BiTE) Blinatumomab (hereto referring to as BiTE), Jurkat and Raji cells interact to form an immune synapse, leading to PD-1/PD-L1 interaction and PD-1 phosphorylation.

Using this setup, we observed that in wild-type PD-1 cells, NT7 robustly co-localized with PD-1 and PD-L1 upon immune synapse formation, indicating ligand-induced phosphorylation of PD-1 **(Fig. 2A, 2B)**. Time-lapse live-cell imaging revealed that phosphorylation occurred rapidly, with NT7-YFP signal at the immune synapse increasing within ∼30–40 seconds and stabilizing thereafter **(Fig. 2C)**. In contrast, NT7 was not recruited to the immune synapse in PD-1 (Y223F/Y248F)-expressing cells, further demonstrating that NT7 specifically recognizes phosphorylated PD-1 **(Fig. 2A, 2B)**. Unlike NT7, the SHP2 tandem SH2 domain (a fusion of N-SH2 and C-SH2) showed minimal co-localization with PD-1 at immune synapses under the same conditions, highlighting that engineered sensors have an even higher sensitivity than native phosphobinding domain even in its tandem form **(Fig. S3C)**. Furthermore, addition of 10 μΜ LCK inhibitor reduced NT7 binding, validating NT7 binding is phosphorylation dependent and consistent with previous reports that LCK phosphorylates PD-1 **(Fig. 2D and E)**.

**Figure 2.**
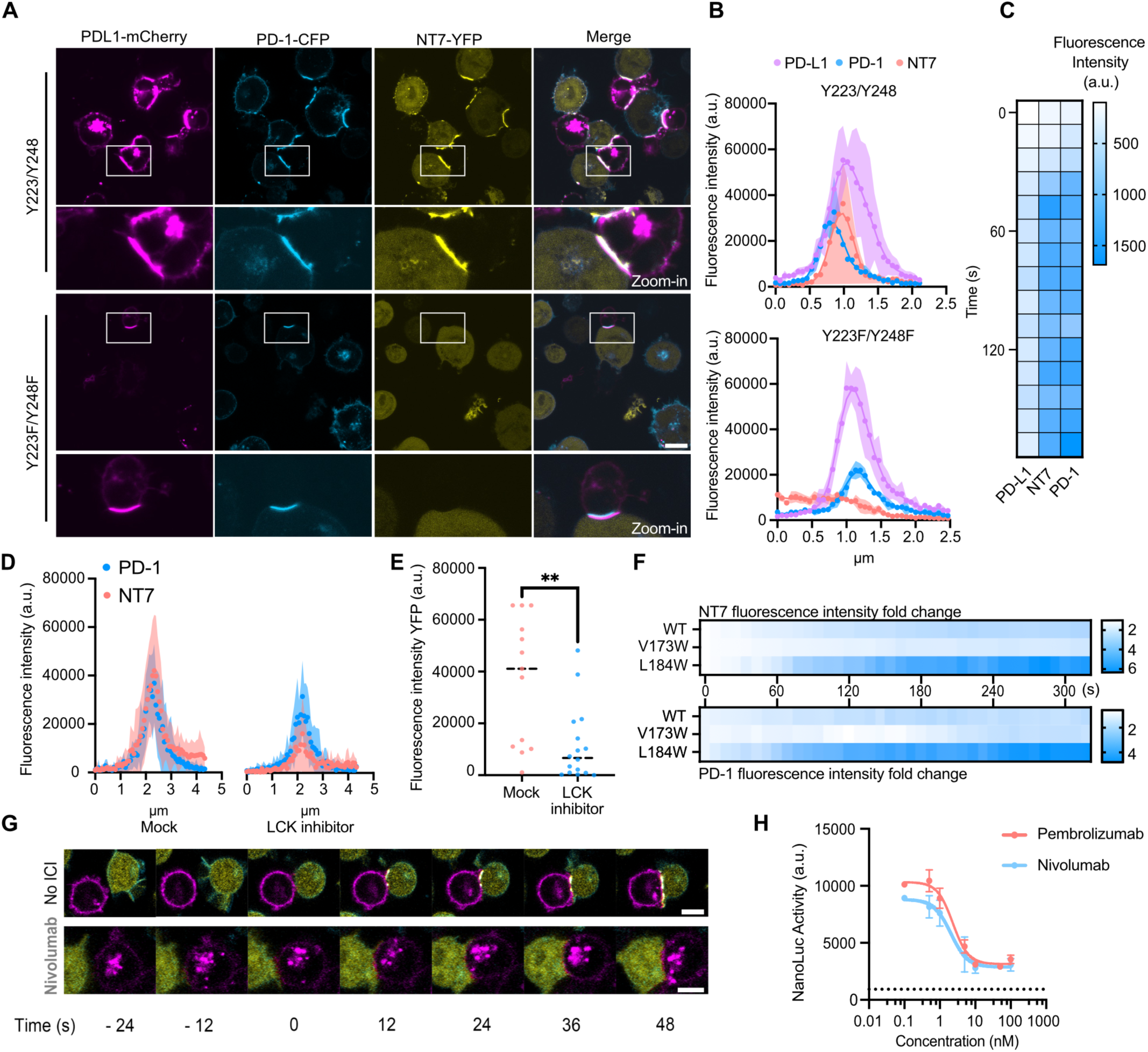
Live-cell Sphyder sensors reveal dynamic PD-1 phosphorylation and regulation by dimerization. **(A)** Representative images of Jurkat T cells expressing wild-type PD-1 (Y223/Y248, top 2 rows) or phosphorylation-deficient PD-1 mutant (Y223F/Y248F, bottom row), co-cultured with PD-L1-mCherry⁺ Raji cells. Wild-type PD-1 shows strong recruitment of the pY248-specific NT7 binder (yellow) to the cell membrane. No NT7 recruitment was observed in cells expressing the double mutant receptor. Scale bars, 10 μm. **(B)** Fluorescence intensity measurements were made across immune synapses, immune synapse was first defined by cell-cell junction, a line segment was drawn across T cell and Raji cell, fluorescence intensity was then determined along the line segment. Comparison of spatial distribution of PD-L1-mCherry, PD-1-CFP, and Sphyder-NT7 binder fluorescence in cells expressing wild-type (top) or mutant (bottom) PD-1. Results represent data from n = 6 synapses. **(C)** Time-resolved heatmap quantification of mean fluorescence intensity for PD-L1, NT7, and PD-1 over time for region of interest, region of interest was drawn as a square to capture the entire interface between T cells and Raji cells. Results represent data from n = 6 synapses. **(D)** Fluorescence intensity profile across the immune synapse in Jurkat cells treated with Lck inhibitor (right) and mock (left). **(E)** Quantification of the NT7 fluorescence intensity at the immune synapse in Jurkat T cells treated with LCK inhibitor compared to mock-treated cells. LCK inhibition significantly reduces PD-1 phosphorylation. **P = 0.0019, unpaired t test, data represent quantification of n = 13 (Mock), n = 16 (LCK inhibitor). **(F)** Time-resolved heatmaps showing fold change in fluorescence intensity for the NT7 (top) and PD-1 (bottom) in cells expressing wild-type PD-1 or transmembrane mutants V173W and L184W. The L184W mutation promotes dimerization and V173W decreases monomerization. Data represents n = 3 synapse. **(G)** Time-lapse confocal imaging of PD-1-CFP⁺ NT7-YFP⁺ Jurkat cells with PD-L1-mCherry⁺ Raji cells showing dynamic immune synapse formation and PD-1 activity. Top row: untreated cells exhibit rapid PD-1 phosphorylation and NT7 recruitment upon synapse formation. Bottom row: Nivolumab pretreatment prevents NT7 recruitment despite normal synapse formation. Time stamps indicate seconds relative to immune synapse formation (0 s). **(H)** Dose-response curves of NanoLuc luciferase activity in PD-1 Split-NanoLuc Jurkat cells treated with varying concentrations of pembrolizumab or nivolumab. Dash line indicates luciferase signal of Split-NanoLuc cells alone. Both ICIs reduce PD-1 phosphorylation signals in a dose-dependent manner. Data represents means ± s.d. from n = 2 independent experiments.

Traditionally, PD-1 has been viewed as a monomeric receptor. More recent studies using Förster resonance energy transfer (FRET) and T cell activity assays, however, indicate that PD-1 can dimerize through interactions within its transmembrane (TM) domain, and that the dimeric form may more potently inhibit T cell signaling compared with the monomeric state (*3*, *19*). Within the TM domain, the L184W mutation promotes PD-1 dimerization, whereas the V173W mutation reduces it. These findings implicate an important role for PD-1 dimerization in receptor activity, but it remains unclear whether PD-1 phosphorylation is directly regulated by the dimeric versus monomeric states, or whether its effects instead arise from enhanced avidity and effector recruitment rather than direct control of phosphorylation.

To investigate the role of dimerization in affecting PD-1 phosphorylation, we expressed PD-1 L184W and V173W mutants fused to CFP in Jurkat cells and co-culture with Raji cells. Confocal imaging revealed that the L184W mutant exhibited a threefold increase in maximal NT7 binding signal compared to wild-type PD-1, whereas the V173W mutant showed a ∼1.5-fold reduction in signal intensity (**Fig. 2F**). The kinetics of phosphorylation of the L184W mutant were comparable to wild type (**Fig. S3D**). These findings indicate that dimerization enhances PD-1 phosphorylation, providing a mechanistic basis for how dimerization strengthens downstream signaling and promotes T cell suppression.

To directly visualize these dynamic events, we performed time-lapse imaging of Jurkat–Raji cell co-cultures (**Movies S1 and S2**). These experiments captured the dynamics of immune synapse formation, PD-1 activation kinetics, and cellular responses to immune checkpoint inhibitors (ICIs). Immune synapses formed in both untreated and ICI-treated cells; however, NT7 recruitment to the synapse was observed only in untreated cells. In cells pretreated with the anti–PD-1 antibody Nivolumab, synapses still formed but NT7 recruitment was abolished, indicating effective blockade of PD-1 phosphorylation (**Fig. 2G**).

Building on these imaging studies, we next developed a high-throughput PD-1 activation assay by engineering split-luciferase biosensors. PD-1 and NT7 were fused to complementary NanoLuc fragments (LgBiT and SmBiT, respectively), such that PD-1 phosphorylation brings the fragments into proximity, reconstituting NanoLuc and generating a luminescence signal upon substrate addition (**Fig. S4A**). Split-NanoLuc Jurkat cells showed low basal luminescence but significantly increased signal when co-cultured with PD-L1⁺ Raji cells and Blinatumomab (**Fig. S4B**). This increase was markedly reduced when PD-1 tyrosine residues Y223 and Y248 were mutated, confirming phosphorylation-dependent NT7 binding (**Fig. S4C**). Several additional Sphyder binders also distinguished wild-type from tyrosine-mutant PD-1 receptors (**Fig. S4C**).

We used the Split-NanoLuc Jurkat cells to assess responses to PD-1/PD-L1 ICIs. Addition of Nivolumab or Atezolizumab (100 nM) to Jurkat/Raji co-cultures treated with BiTE rapidly decreased luminescence, reflecting ICI-mediated PD-1 dephosphorylation, while a control anti-EGFR antibody had no effect (**Fig. S4D**). The assay also allowed potency measurements, yielding IC_50_ values for Pembrolizumab (2.3 nM) and Nivolumab (2.0 nM) consistent with previous reports (**Fig. 2H**) (*24*). We also tested Peresolimab, a clinical-stage PD-1 agonist antibody, which induced a dose-dependent luminescence increase in co-cultures of PD-1 Split-NanoLuc Jurkat cells and FcRγ⁺, PD-L1⁻ THP-1 cells (**Fig. S4E**). Interestingly, this effect was absent in PD-L1⁺ Raji cells lacking FcRγ, suggesting that FcRγ engagement is necessary for Peresolimab’s activity (**Fig. S4F**). At higher concentrations (10–100 nM), the signal declined, showing a hook effect typical of bispecific molecules (*25*). These findings are consistent with reports that Peresolimab enhances PD-1 signaling by promoting PD-1–TCR co-localization through FcRγ-dependent interactions (*26*), and highlight the utility of PD-1/NT7 Split-NanoLuc cells as a high-throughput and quantitative platform for characterizing or screening PD-1 inhibitors and activators.

### Expanding the Sphyder designs to LAT pY161

We next interrogated whether this platform could be expanded to other phosphotyrosine motifs. We focused on LAT, an adaptor protein that plays a critical role in TCR signaling by assembling multiprotein complexes upon phosphorylation at site Y161 (*5*). As with PD-1, native SH2 domains bind LAT phosphotyrosine motifs with low affinity and modest specificity, limiting their utility as sensors or engineering building blocks (*16*). To identify suitable SH2 scaffolds for LAT targeting, we first tested the binding of dual SH2 domains from PLCγ1, which interacts with phosphorylated LAT to promote TCR signaling **(Fig. S5A)** (*27*). Using yeast surface display, we evaluated both the PLCγ1 N-terminal SH2 domain in its wild-type form and a variant carrying previously described “superbinder” mutations intended to enhance phosphotyrosine binding **(Fig. S5B)** (*10*). While the wild-type PLCγ1 SH2 domain showed limited binding to LAT pY161, the superbinder variant exhibited stronger binding affinity and was thus selected as the SH2 module for Sphyder binder engineering **(Fig. S5B)**.

Analogous to our PD-1 strategy, we performed circular permutation on the PLCγ1 superbinder SH2 domain to generate new N- and C-termini suitable for fusion to a Nb library. We constructed a yeast surface display library encoding the permuted PLCγ1 SH2 domain fused via a flexible glycine-serine linker to randomized Nbs, achieving a library diversity of ∼2 × 10^6^ unique variants **(Fig. 3A)**. This library was screened using gradually decreasing concentrations of biotinylated LAT pY161 peptide, with additional negative selection steps to remove clones binding unphosphorylated LAT peptides. Successive rounds of FACS led to clear enrichment of clones exhibiting higher fluorescence intensity, indicating enrichment of high-affinity clones for phosphorylated LAT **(Fig. S5C)**.

**Figure 3.**
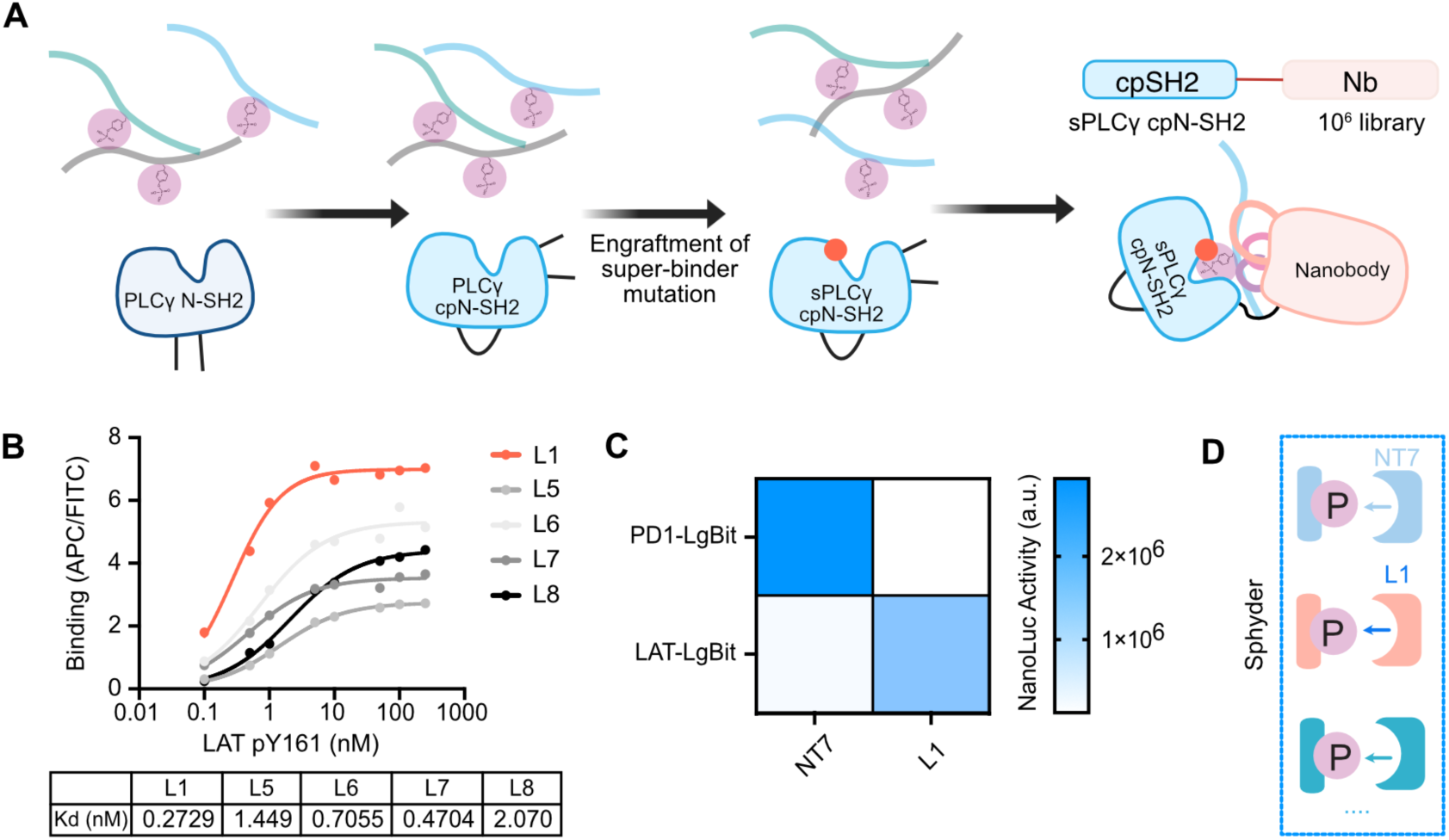
Generalization of the Sphyder platform to generate binders targeting LAT pY161. **(A)** Schematic of the Sphyder binder design strategy for LAT pY161. A circularly permuted PLCγ1 N-terminal SH2 domain (blue) is used as the primary phosphotyrosine recognition module, enabling fusion of a nanobody library for enhanced specificity and affinity toward LAT pY161. **(B)** Yeast surface titration curves showing binding of selected Sphyder LAT binders (L1, L5, L6, L7, L8) to biotinylated LAT pY161 peptide. The dissociation constants (Kd) are indicated in the table below, ranging from sub-nanomolar to low nanomolar affinities, demonstrating successful engineering of high-affinity LAT phosphosite binders. **(C)** Split NanoLuc luciferase assay results confirming specificity of Sphyder binders for LAT pY161. Calibration bars indicate luminescence signal from combinations of PD-1-LgBiT or LAT-LgBiT fused constructs with either Sphyder PD-1-SmBiT binder or Sphyder LAT-SmBiT binder. Strong luminescence is observed only for cognate binder–target pairs, confirming the specificity of Sphyder LAT binders and minimal cross-reactivity with PD-1 pY248. **(D)** Schematics of specific recognition of immune phosphotyrosine signaling sequence by Sphyder binders.

NGS of the enriched pools identified multiple unique binder sequences, among which the top five enriched candidates (L1, L5, L6, L7, and L8) were synthesized and characterized individually. Yeast titration assays revealed that these Sphyder LAT binders exhibited low nanomolar dissociation constants (Kd ranging from ∼0.27 to ∼2 nM), significantly surpassing the affinity of the PLCγ1 SH2 domain alone **(Fig. 3B)**. Specificity profiling against a panel of unrelated phosphopeptides, including PD-1 pY248, confirmed that these LAT binders were highly selective for the LAT pY161 epitope **(Fig. S5D)**.

To demonstrate functionality in living cells, we fused the LAT binder L1 to a fluorescent protein domain and expressed it in Jurkat T cells. Confocal microscopy of the Jurkat cells co-cultured with Raji cells activated with BiTE revealed strong co-localization of the Sphyder-LAT binder with LAT-mCherry at the immune synapse, confirming specific recognition of phosphorylated LAT in live cells **(Fig. S5E**, upper panel without tumor cells, lower panel with tumor cells). Furthermore, we validated the binder’s specificity and phospho-dependence using a split NanoLuc luciferase assay, in which LAT-LgBiT reconstituted luciferase activity only when paired with its cognate Sphyder LAT binder and not with PD-1-specific Sphyder binders **(Fig. 3C)**. In Jurkat cells, expression of LAT-LgBit fusion and Sphyder LAT binder-SmBit fusion reported LAT phosphorylation in response to CD3/CD28 co-stimulation **(Fig. S5F)**. These results demonstrate that the Sphyder platform can be extended beyond PD-1 to generate highly specific orthogonal binders for additional phospho-motifs **(Fig. 3D)**.

### Synthetic immune signaling module for programming phosphorylation pathways

Anti-PD-1 ICIs disrupt PD-1/PD-L1 interactions to restore T cell function. However, this approach only blocks extracellular ligand engagement, leaving ligand-independent signaling events and compensatory pathways unaffected, and therefore may not achieve complete signal suppression (*28*, *29*). In CAR-T therapy, PD-1 signaling contributes to exhaustion, but PD-1 blockade or knockout strategies have yielded mixed outcomes in restoring T cell activity (*30–32*). We hypothesized that synthetic signaling modules could be engineered to sense phosphorylation events and convert them into defined outputs using Sphyder phosphotyrosine–synthetic binder pairs, providing an intracellular strategy to more effectively rewire PD-1 signaling outcomes. Unlike endogenous binders that recognize tandem motifs with limited specificity, Sphyder binders such as NT7 selectively recognize defined phosphotyrosine sequences. This enables minimal phospho-motifs to be paired with their cognate binders to construct synthetic signaling pathways with programmable outputs (**Fig. 4A**).

**Figure 4.**
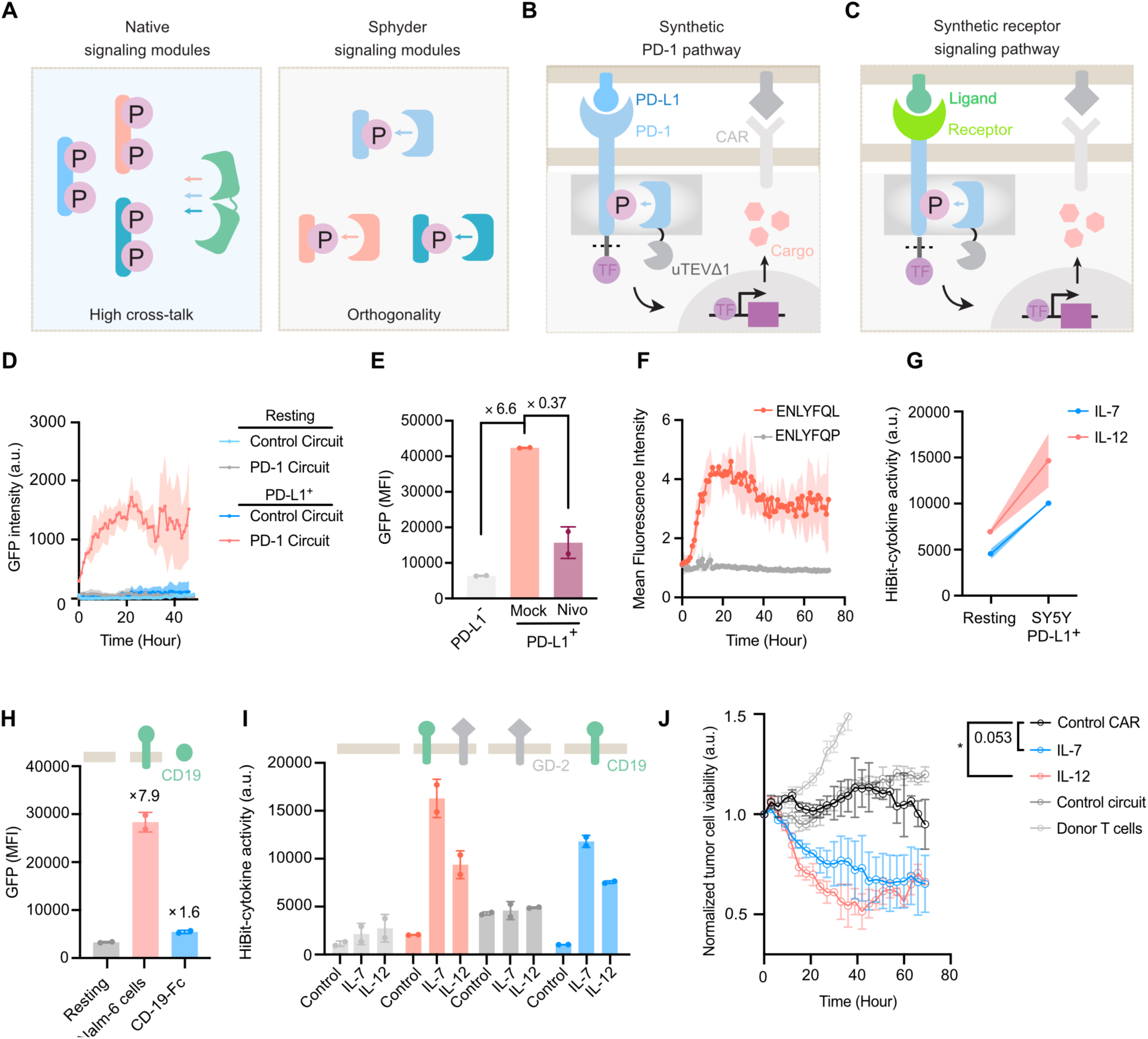
Synthetic rewiring of PD-1 signaling into immunoactivating outputs using Sphyder circuits. **(A)** Comparison of native versus Sphyder signaling modules. Native phosphotyrosine signaling modules often bind tandem motifs with overlapping specificity, resulting in signaling cross-talk. In contrast, Sphyder generates orthogonal phosphotyrosine–synthetic binder pairs in which each binder recognizes a defined single phosphotyrosine motif with high affinity and specificity. This orthogonality enables the construction of synthetic signaling pathways and receptors with customizable input–output properties. **(B)** Engineered PD-1 pathway using Sphyder binders. Phosphorylated PD-1 recruits synthetic modules that convert inhibitory signaling into immune activation via inducible gene expression. NT7 fused to TEV protease enables phosphorylation-dependent cleavage of transcription factors (e.g., Gal4, tTA) tethered to PD-1 via TEVs, initiating gene expression. **(C)** Schematics of using Sphyder binders to reprogram receptor signaling. A customized antigen recognition domain replaced the ectodomain of PD-1 while retaining the intracellular domain of PD-1. Similar to synthetic PD-1 signaling, a transcription factor would be released upon phosphorylation to drive antigen-dependent response. **(D)** Time kinetics of TRE3G driven GFP expression in PD-1 circuit, GFP was upregulated in PD1-TEVs-tTA and NT7-uTEVΔ1 expressing primary T cells co-cultured with PD-L1^+^ Raji cells at early time points and with sustained level for 2 days compared to non co-culture or cells only transduced with TRE3G::GFP. **(E)** GFP expression quantification in Jurkat T cells expressing PD-1 Sphyder circuits. Cells were co-cultured with PD-L1- or PD-L1⁺ Raji cells in absence of presence of 100 nM PD-1 inhibitor Nivolumab (Nivo). PD-1 Sphyder circuit-expressing cells show strong PD-L1-induced GFP expression, while cells exposed to PD-L1⁺ in presence of Nivolumab exhibit reduced GFP levels. **(F)** Time-course analysis of GFP expression in the Sphyder-circuit using IncuCyte. GFP expression is rapidly induced following co-culture with PD-L1^+^ Raji cells and TEV-mediated cleavage. The TEV-uncleavable variant (ENLYFQP) showed minimal activation. Data represent mean ± s.d. from 2 replicates, each well were imaged 9 images for averaging and quantification. **(G)** NanoLuc luciferase assay quantifying HiBiT-tagged IL-7 (Blue) and IL-12 (Red) secretion in media in response to PD-1 activation by SY5Y PD-L1^+^ cells. Red: IL-12, blue: IL-7. **(H)** 7.9-fold induction of GFP was observed in cells expressing FMC63-replaced receptor cultured with CD19^+^ Raji cells, or 1.6-fold induction when co-cultured with a soluble bivalent CD19-Fc fusion protein. **(I)** A CD-19 driven Sphyder circuit in GD2-HA CAR-T cells showed CD-19 dependent cytokine secretion, co-culture with a GD-2 positive but CD-19 negative tumor cell line did not induce cytokine secretion from engineered T cells. Data represent mean ± s.d. from 2 replicates. **(J)** Time-course quantification of tumor cell killing by primary CAR-T cells expressing PD-1 circuits driving IL-7 and IL-12 expression. Sphyder-based transcriptional circuits enhance tumor clearance relative to control CAR-T cells. Two-way ANOVA, *P=0.0131. Data represent mean ± s.d. from 2 replicates.

We developed two Sphyder-based cellular pathways that rewire PD-1 phosphorylation into either (1) kinase activation (**Fig. S6A**) or (2) inducible gene expression (**Fig. 4B**). For driving kinase activity, we fused the interdomain B and kinase domain of ZAP70 (Zeta-chain-associated protein kinase 70), an enzyme downstream TCR signaling for T cell activation to NT7 (*33*). Confocal imaging confirmed phospho-dependent recruitment of NT7-ZAP70 to the plasma membrane in Jurkat cells expressing wild-type PD-1 but not phosphorylation-deficient mutants (**Fig. S6A**) when co-culture with PD-L1^+^ Raji cells and activated with BiTE. Functionally, GD2-targeting HA-28z CAR-T cells expressing NT7-ZAP70 exhibited enhanced tumor killing after repeated challenge with GD2⁺ PD-L1⁺ SY5Y cells, demonstrating that synthetic phospho-sensing modules can redirect kinases to inhibitory receptors and enhance effector function (**Figs. S6B– C**).

In a second design, we built a synthetic protease-based transcriptional circuit that converts PD-1 phosphorylation into designed transcriptional programs. NT7 was fused to the Tobacco etch virus (TEV) protease to generate a phosphorylation-dependent protease module (NT7-TEV) (**Fig. 4B–C**). The PD-1 intracellular domain was fused to a TEV substrate sequence (TEVs) and a synthetic transcription factor (tTA or Gal4), creating PD-1-TEVs-tTA or PD-1-TEVs-Gal4 constructs. Upon PD-1 phosphorylation, NT7-TEV was recruited to PD-1 and cleaved the TEVs fusion, releasing tTA or Gal4 to translocate into the nucleus and activate downstream genes (e.g., GFP). In LCK-transfected HEK293T cells expressing PD-1-TEVs-tTA, NT7-TEV localized to the membrane and released tTA into the nucleus (**Fig. S6D, top**). In contrast, cells expressing a CD28-TEVs-tTA construct showed minimal NT7-TEV recruitment or nuclear release, confirming PD-1-specific activation (**Fig. S6D, bottom**). In human primary T cells, the circuit induced GFP expression in a PD-1-phosphorylation-dependent manner **(Fig. S6E)**. Live-cell imaging revealed GFP induction within 1 hour of PD-1 activation, with sustained expression for over 48 hours (**Fig. 4D**). GFP induction was reduced by Nivolumab treatment (**Fig. 4E**) and abolished when the TEVs was replaced with an uncleavable TEV substrate (ENLYFQ/L –> EGLYFQ/P) (**Fig. 4F**), confirming that circuit activation required TEV-mediated cleavage.

To adapt this circuit for therapeutic applications, we tested whether it could drive local cytokine expression. We focused on IL-12 and IL-7 because of their complementary roles in anti-tumor immunity. IL-12 enhances cytotoxic T and NK cell activity, IFN-γ secretion, and Th1 polarization and its systemic administration is limited by toxicity (*34*, *35*). IL-7, on the other hand, facilitates T cell survival, proliferation, and memory formation (*36*). In both Jurkat and primary human T cells expressing the NT7-based Sphyder circuit, PD-1 activation induced by PD-L1–expressing Raji cells triggered expression of IL-7 and an engineered full-length IL-12, a single-chain variant that fuses the p35 and p40 subunits of the IL-12 complex (*37*, *38*) (hereafter referred to as IL-12) (**Fig. S6F**, **Fig. 4G**). These results demonstrate the successful construction of a synthetic PD-1 signaling pathway that enables PD-1 activation–gated cytokine expression using Sphyder.

In the tumor microenvironment, T cells encounter diverse extracellular cues that shape their function and fate. To test whether the Sphyder signaling module could be leveraged to construct synthetic receptors with customizable input–output properties, we studied whether the extracellular domain of PD-1 could be modularly replaced to enable programmable sensing of non-canonical inputs and rewiring of those signals into defined intracellular responses. Specifically, we substituted the PD-1 ectodomain with FMC63, a single-chain antibody fragment that recognizes CD, or replaced the intracellular signaling motif of an FMC63-based CAR with the 14–amino acid segment surrounding PD-1 pY248 to generate a CD19-targeting PD1-ITSM chimeric receptor (**Fig. 4C**). When paired with a TEV–tTA Sphyder circuit, both chimeric receptors enabled CD19-dependent induction of GFP expression in Jurkat cells (**Fig. 4H, Fig. S7A**). Building on this design, we expressed this circuit in primary CAR-T cells to drive IL-7 or IL-12 expression. Co-culture with CD19⁺ Nalm6 cells resulted in sustained expression of both cytokines, demonstrating the capacity of Sphyder-based receptors to convert specific antigen recognition into therapeutic transcriptional programs (**Fig. S7B**). Additionally, we expressed the CD19-targeting PD1-ITSM chimeric receptor in GD2-targeting HA-28z CAR-T cells. Cytokine secretion was observed only when the T cells were co-cultured with CD19⁺ Nalm6 or dual-antigen CD19⁺/GD2⁺ cells, but not with CD19⁻ GD2⁺ cells, confirming that cytokine expression was specifically gated by the CD19-targeting Sphyder chimeric receptor rather than the GD2-targeting CAR signaling (**Fig. 4I**).

To further evaluate the therapeutic potential of cytokine-expressing Sphyder circuits, we transduced PD-1-TEVs-tTA, NT7-TEV, and the cytokine gene circuit into human primary GD2-targeting HA-28z CAR-T cells. PD-1-activation induced IL-7 or IL-12 expression enhanced tumor killing in co-culture with PD-L1⁺ neuroblastoma SY5Y cells compared to control CAR-T cells lacking the circuit (**Fig. 4J**). These findings demonstrate that Sphyder-based circuits can rewire inhibitory checkpoint signals into tailored intracellular programs in primary T cells and may provide a strategy to improve the efficacy and persistence of CAR-T therapies.

### Ivonescimab induces VEGF-dependent PD-1 degradation

Anti–PD-1 ICIs remain a cornerstone of cancer immunotherapy. Recently, bispecific antibodies that simultaneously target PD-1 and VEGF have gained considerable attention. Ivonescimab, a PD-1/VEGF bispecific antibody, is constructed by fusing a PD-1–blocking scFv onto the C-terminus of the IgG scaffold of bevacizumab, a VEGF-neutralizing antibody. In the global HARMONi-1 trial, Ivonescimab combined with chemotherapy improved outcomes for patients with EGFR-mutated non–small cell lung cancer (NSCLC) who had progressed on third-generation tyrosine kinase inhibitors, extending median progression-free survival (PFS) to ∼6.8 months versus ∼4.4 months for chemotherapy alone (hazard ratio 0.52; 95% CI 0.41-0.66; p < 0.00001) (*39*, *40*), with encouraging but statistically borderline overall survival trends (p = 0.057) (WCLC 2025). In parallel, the HARMONi-2 trial compared Ivonescimab monotherapy with pembrolizumab in first-line PD-L1⁺ NSCLC, showing a striking median PFS of ∼11.1 months versus ∼5.8 months (HR ∼0.51, p < 0.0001) (*41*). While overall survival data from HARMONi-2 are not yet available, these results have positioned Ivonescimab as one of the most clinically advanced bispecific checkpoint inhibitors and generated strong interest in understanding whether anti-PD-1/VEGF antibodies may become the next generation of immunotherapy.

Dual pathway blockade of PD-1 and VEGF by combination of PD-1 and VEGF antibodies has shown mixed clinical outcomes (*42*, *43*). However, Ivonescimab appears to deliver benefits beyond what has historically been seen with combinations of separate PD-1 and VEGF antibodies. This raises important mechanistic questions: does Ivonescimab engage PD-1 signaling in a fundamentally different way from other PD-1/PD-L1 inhibitors, or does its bispecific architecture confer unique activity not achieved by combining monospecific PD-1 and VEGF antibodies.

We applied Sphyder PD-1 sensors to characterize the effects of Ivonescimab in living cells. We first examined the immediate effects of Ivonescimab on PD-1 signaling by analyzing Jurkat–Raji cocultures 30 minutes after treatment with the antibodies in the presence of BiTE. In untreated cells, NT7 strongly localized to the immune synapse, indicating PD-1 phosphorylation (**Fig. 5A, see also Fig. 2A**). When Ivonescimab was added in the absence of VEGF, NT7 recruitment at synapses was reduced in subsets of synapses (**Fig. 5B**), consistent with partial blockade of PD-1 phosphorylation through competitive inhibition of PD-L1 binding, similar to conventional ICIs. At 10 nM, Ivonescimab showed weaker inhibition than Nivolumab, which at the same concentration completely abolished NT7 recruitment (**Fig. 2G**).

**Figure 5.**
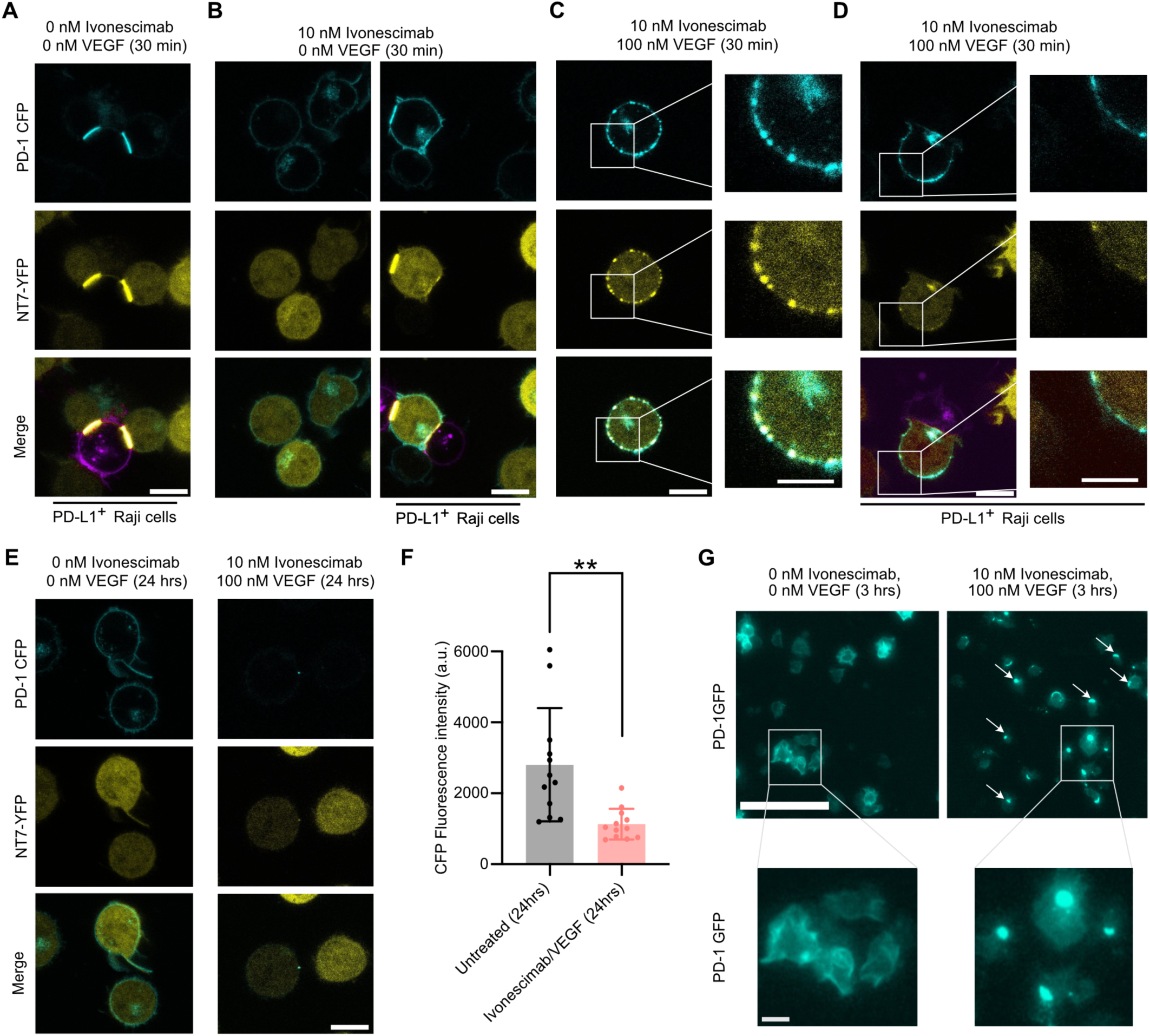
Ivonescimab induces VEGF-dependent PD-1 clustering, phosphorylation, and receptor degradation. (A-E) Representative confocal images of PD-1-CFP⁺ NT7-YFP⁺ Jurkat cells treated with 10 nM Ivonescimab in the presence of 0 or 100 nM VEGF at 30 min or 24 hrs. Scale bar, 10 μm. Scale bar for zoom-in images, 5 µm. **(A)** Cells without Ivonescimab treatment co-cultured with PD-L1⁺ Raji cells showed strong PD-1 phosphorylation and NT7-YFP recruitment at the immune synapse. **(B)** Under 0 nM VEGF conditions, 30 minutes of Ivonescimab treatment partially inhibited PD-1 activation, as indicated by reduced NT7-YFP recruitment in one of the two immune synapses. **(C)** With 100 nM VEGF, 30 minutes of Ivonescimab treatment induced prominent PD-1 clustering at the cell surface and NT7-YFP recruitment, consistent with Y248 phosphorylation. **(D)** Jurkat cells co-cultured with PD-L1⁺ Raji cells and treated with Ivonescimab and VEGF (30 min) showed partial inhibition of PD-1 phosphorylation at the immune synapse; however, membrane clustering of PD-1 with NT7 recruitment was still evident. **(E)** With 100 nM VEGF, 24 hours of Ivonescimab treatment induced substantial loss of surface PD-1-CFP signal (Right panel), compared to untreated cells (Left panel). **(F)** Quantification of surface PD-1-CFP fluorescence intensity in untreated versus Ivonescimab-treated cells showed significant reduction of PD-1 on the plasma membrane at 24 hrs. Student t-test, P = 0.0107. **(G)** IncuCyte live-cell imaging of Jurkat cells expressing PD-1-GFP treated with Ivonescimab and VEGF for 3 hours showed redistribution of PD-1 into intracellular puncta. White arrows highlight clustering and internalizing puncta. Scale bar, 100 μm, scale bar in the zoom in panel, 10 μm.

Strikingly, the addition of VEGF to Ivonescimab drove prominent PD-1 clustering across the plasma membrane, accompanied by strong NT7 recruitment (**Fig. 5C**). Clusters were observed both in both presence and absence of PD-L1+ Raji cells, indicating that Ivonescimab induces PD-1 clustering independently of target cell engagement (**Fig. 5D**).

To understand the long-term consequences of this clustering, we monitored PD-1 surface expression over extended treatment periods. In PD-1–GFP⁺ Jurkat cells treated with Ivonescimab and VEGF for 24 hours, PD-1 fluorescence at the plasma membrane was markedly reduced compared with untreated controls (**Fig. 5E**), and quantification confirmed a significant reduction in surface PD-1 levels (p = 0.0107, **Fig. 5F**). Importantly, redistribution of PD-1 into intracellular puncta was already visible within 3 hours of treatment and occurs almost universally across all cells (**Fig. 5G and Fig. S8A)**, indicating that clustering initiates rapid receptor internalization. These findings revealed that Ivonescimab induces VEGF-dependent PD-1 clustering followed by receptor degradation.

### Ivonescimab exhibits transient dual agonist/antagonist-like activity

We further characterized the time-dependent activity of Ivonescimab. Quantification of NT7-YFP and PD-1-CFP fluorescence intensity along the plasma membrane using the Oval Profile tool (*44*) confirmed that Ivonescimab induced pronounced PD-1 clustering accompanied by NT7 co-localization in the presence of VEGF, confirming phosphorylation at the clustering sites (**Fig. 6A–B**). To measure this effect quantitatively, we used Split-NanoLuc Jurkat cells. At 30 minutes, Ivonescimab plus VEGF triggered a VEGF dose-dependent luminescence increase (**Fig. 6C**), indicating enhanced PD-1 phosphorylation both in the absence and presence of PD-L1⁺ Raji cells. This phosphorylation was abolished by the Src-family kinase inhibitor Dasatinib (**Fig. S8B**), indicating that the signal reflected LCK-mediated PD-1 phosphorylation. In contrast, in the absence of VEGF, Ivonescimab behaved as a conventional antagonist, reducing PD-1 phosphorylation with an IC_50_ of ∼10 nM, approximately fivefold weaker than pembrolizumab or nivolumab (IC_50_ ∼2 nM, **Fig. 2H**, measured at 30 min). VEGF progressively attenuated this inhibitory effect: at 40 nM VEGF the IC_50_ shifted more than tenfold higher (**Fig. S8D**), and at 80 nM VEGF Ivonescimab no longer inhibited PD-1 phosphorylation (**Fig. 6D, red curve**).

**Figure 6.**
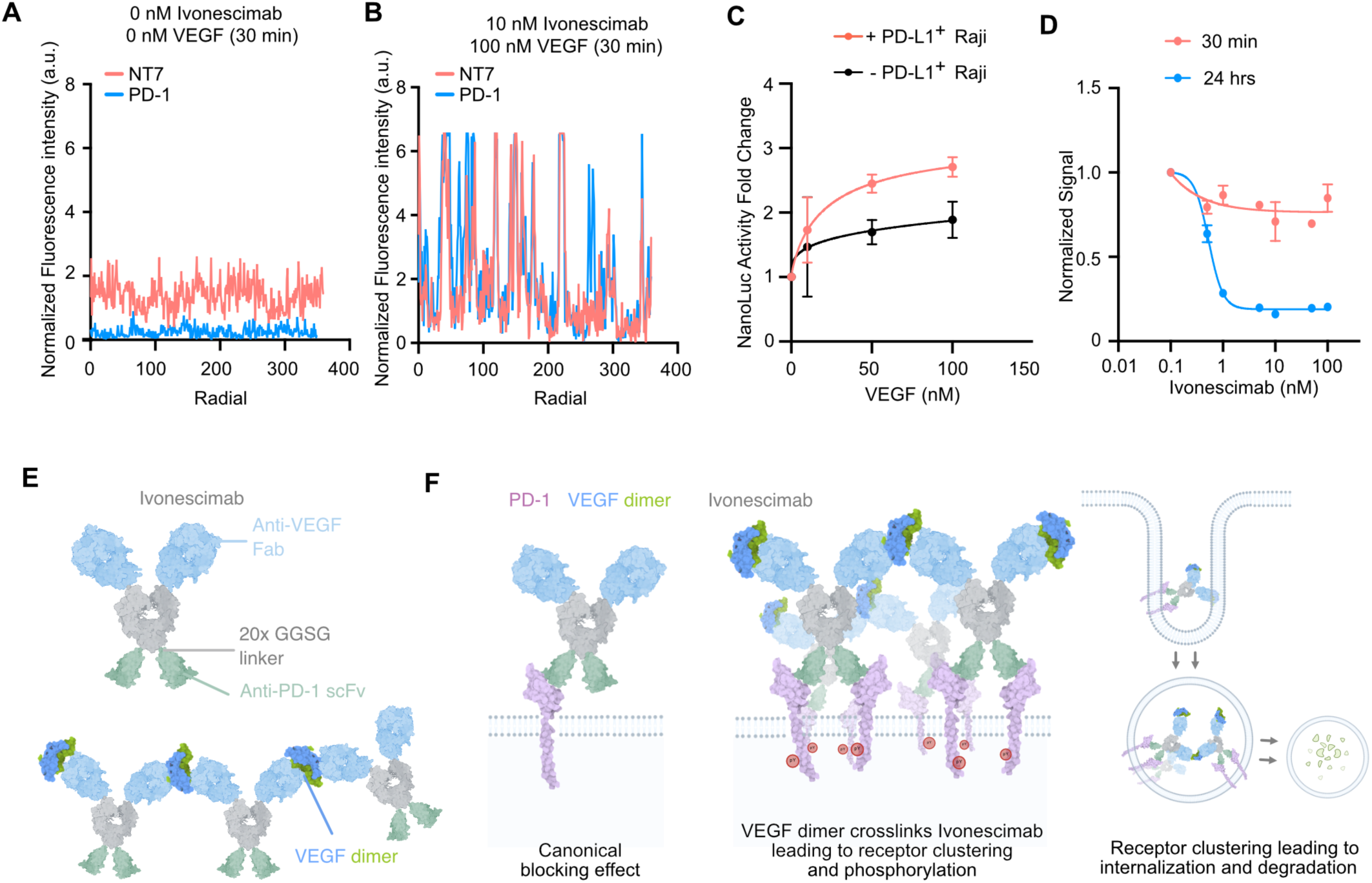
Model of Ivonescimab activity. (A-B) Quantification of NT7-YFP and PD-1-CFP fluorescence intensity along the plasma membrane of Jurkat cells untreated (A) or treated (B) with Ivonescimab plus VEGF. Untreated cells showed uniform membrane distribution without clustering, whereas Ivonescimab plus VEGF induced overlapping peaks of NT7 and PD-1 signals. Fluorescence intensity was measured using the Oval Profile tool, with radial distance corresponding to the membrane circumference. Data are shown from one representative cell for each condition. **(C)** Normalized luciferase activity showing a dose-dependent VEGF-mediated enhancement of PD-1 phosphorylation after 30 min treatment with 10 nM Ivonescimab in Split-NanoLuc Jurkat cells. Data are shown for co-cultures with or without PD-L1⁺ Raji cells in the presence of 1 nM BiTE. Values represent mean ± standard deviation from n = 2 replicates. **(D)** Normalized NanoLuc luciferase activity in Split-NanoLuc Jurkat cells co-cultured with PD-L1⁺ Raji cells plus 1 nM BiTE, treated with increasing doses of Ivonescimab for 30 min or 24 h. VEGF concentration was 80 nM for the 30 min condition and 100 nM for the 24 h condition. At 30 min, Ivonescimab in the presence of VEGF acted as a neutral binder, whereas at 24 h it functioned as a potent antagonist with a sub-nM IC_50_ Data for additional VEGF concentrations are provided in Fig. S8. Values represent mean ± standard deviation from n = 2 replicates. **(E)** Schematic of Ivonescimab structure, showing anti-VEGF Fab regions (blue), anti–PD-1 scFv domains (green), linker between the Fc and anti-PD-1 scFv domain (grey), and VEGF dimer (blue/green) mediating crosslinking with Ivonescimab. **(F)** Proposed model of Ivonescimab mechanism of action. Left: In the absence of VEGF, Ivonescimab acts as a conventional PD-1 blocking antibody and blocks PD-L1 binding. Middle: in the presence of VEGF dimers, Ivonescimab is crosslinked into higher-order complexes, leading to PD-1 clustering and phosphorylation. Right: clustered PD-1 is subsequently internalized and trafficked to lysosomes for degradation.

We then examined how these early signaling effects evolved over time. After 24 hours of treatment, Ivonescimab in the presence of VEGF acted as a potent antagonist, suppressing PD-1 phosphorylation with a sub-nM IC_50_ (**Fig. 6D, blue curve; Fig. S8E–F**). The antagonist activity at 24 h is consistent with live-cell imaging showing that Ivonescimab promotes VEGF-dependent PD-1 internalization and degradation over time (**Figs. 5E–G, S8A**).

Because VEGF is a homodimer (*45*), its engagement with Ivonescimab creates opportunities for multivalent antibody–ligand assemblies that may underlie the observed receptor clustering and internalization activity **(Fig. 6E)**. Our data support a model in which Ivonescimab acts through context-specific, time-dependent mechanisms. In the absence of VEGF, it functions as a conventional antagonist, blocking PD-1 interaction with PD-L1 in a manner similar to other ICIs (**Fig. 6F, left panel**). In the presence of VEGF, dimeric VEGF crosslinks Ivonescimab into higher-order oligomeric complexes through dimer-on-dimer “daisy chain” interactions, driving rapid PD-1 clustering at the cell surface (**Fig. 6E**; **Fig. 6F, middle panel**). The clustered PD-1 is subsequently internalized, removed from the plasma membrane, and degraded intracellularly (**Fig. 6F, right panel**). This VEGF-dependent oligomerization model is consistent with recent biochemical studies reporting large Ivonescimab–VEGF complex formation detected by size-exclusion chromatography (*46*).

In addition to promoting receptor degradation, clustering was accompanied by PD-1 phosphorylation, revealing a transient agonist-like phase before induced receptor clearance. This dual activity resembles recent findings that antagonistic checkpoint antibodies with reduced affinity can paradoxically agonize their targets through clustering mechanisms (*47*). The functional consequences of the transient phase of combined agonistic and antagonistic effects of Ivonescimab for T cell activity remain to be determined.

These results provide mechanistic insights into the activity of this new class of bispecific ICIs and suggest potential avenues for molecular optimization. For instance, VEGF-driven receptor degradation may contribute to the long-term efficacy of anti-PD-1/VEGF bispecific antibodies, as targeted receptor clearance may provide more durable inhibition than conventional neutralizing approaches (*48*). Additionally, because dimer-on-dimer complex formation appears important for activity, parameters such as epitope geometry, domain orientation, and linker length may influence PD-1 clustering, degradation efficiency, and VEGF sensitivity, presenting opportunities to further optimize Ivonescimab or similar antibodies for enhanced activity.

## Discussion

Cells rely on phosphorylation as an essential mechanism for regulating both surface receptor activity and intracellular signaling cascades. Accordingly, therapeutics that influence phosphorylation, such as kinase inhibitors (*49*), monoclonal antibodies targeting receptor tyrosine kinases (*50*), and immune checkpoint inhibitors (*51*), have transformed the treatment of cancer and immune-related diseases. Despite this success, molecular insight into how these therapies modulate receptor activity at the signaling level remains limited, owing to a lack of tools for reporting signaling activation within living cells. Beyond sensing, the ability to reprogram phosphorylation-dependent signaling at defined sites represents a powerful yet largely untapped strategy for therapeutic cell reprogramming. While synthetic biology has begun to explore these opportunities, efforts are often constrained by the lack of sequence specificity needed to achieve targeted outcomes (*52*, *53*).

Our study addresses these needs by introducing a strategy for sensing and rewiring immune signaling through precise phosphosite recognition. Building on prior work developing engineered phosphotyrosine-binding domains, such as our pY-TRAP system (*14*) and the pY-Clamp method (*54*), which established the concept of cooperative, two-domain sandwich architectures for phosphosite recognition, we extend this approach for the first time to immune signaling protein signaling motifs. By leveraging domains from SHP2 and PLCγ1, we created nanomolar-affinity, site-specific synthetic binders targeting PD-1 and LAT motifs. These binders functioned as modular components for both real-time biosensors and synthetic circuits, enabling mechanistic interrogation and programmable rewiring of immune signaling with site specificity.

Although protein design methods have advanced rapidly, strategies for creating protein binders that can distinguish a post-translational modification only within its specific amino acid sequence context remain unavailable (*55*). This challenge likely reflects the difficulty of engineering binding modules to simultaneously recognize both the modification and the surrounding residues. Our results, demonstrating nanomolar-affinity binders with both phospho- and sequence-specificity, suggest that cooperative, multi-domain binding architectures offer a viable design principle to address this challenge. Such designs may be incorporated into future strategies for developing binders to peptides carrying diverse post-translational modifications, including but not limited to phosphorylation. This design principle also resembles molecular glue drug mechanisms (*25*), where a ligand sandwiched between two protein domains can dramatically enhance overall binding affinity, though here we apply the concept in an entirely new context.

The Sphyder biosensor for PD-1 phosphorylation illustrates the utility of this approach for studying receptor regulation. Unlike transcription-based or biochemical readouts of receptor activity (*56*, *57*), these sensors directly report the changes in PD-1 phosphorylation in living cells through rapid fluorescence or luminescence signals, thereby providing spatiotemporal information on receptor regulation. The biosensor detected rapid dephosphorylation induced by PD-1/PD-L1 blocking antibodies, which may reflect displacement of PD-1 from the immune synapse and CD45-mediated dephosphorylation in the presence of ICIs, a mechanism that warrants further study (*58*, *59*). We also observed agonist activity of the clinical antibody Peresolimab in FcR⁺ THP-1 cells but not in FcR⁻ Raji cells, consistent with an FcR engagement mechanism (*60*). Furthermore, the biosensor revealed that receptor dimerization increases PD-1 phosphorylation, consistent with recent findings showing that dimerization affects PD-1 function (*3*, *19*). This observation suggests a potential path for developing PD-1 modulators that regulate its dimerization state to control receptor activity.

Beyond providing insight into both physiological regulation and pharmacological effects, Sphyder enables perturbation and rewiring of immune receptor signaling. In contrast to current strategies that block extracellular PD-1/PD-L1 interaction, our approach modulates signaling from within the cell. Rather than simply inhibiting PD-1 activity, we engineered circuits that redirect its intracellular output toward an immunoactivation or modulatory effect. This design may thus allow modulation of both ligand-dependent and ligand-independent (tonic) PD-1 signaling (*28*) and supports the engineering of immune cells with programmable therapeutic functions. We demonstrated platform modularity at two levels: (i) by engineering binders to both PD-1 and LAT motifs, underscoring target generality, and (ii) by reconfiguring binder–phosphosite modules into customizable input–output circuits. This modularity may enable programmable sensing across many immune receptors and tailoring of responses to specific therapeutic needs.

Our study provides mechanistic insight into the function of anti–PD-1/VEGF bispecific antibodies, a therapeutic class under active clinical evaluation. In addition to mechanisms shared with conventional PD-1 inhibitors that block ligand–receptor interactions, we found that Ivonescimab also induces VEGF-dependent clustering and phosphorylation of PD-1, followed by receptor degradation. We hypothesize that this clustering is driven by oligomerization of Ivonescimab upon VEGF binding, consistent with recent biochemical studies showing that VEGF and Ivonescimab can form various oligomeric complexes at different molar ratios (*46*, *61*). While the clinical implications remain to be determined, these findings provide hypotheses to explain the differentiated activity of these antibodies and suggest potential avenues for further engineering and tailoring of their function.

This mechanism also connects to the emerging field of extracellular targeted protein degradation (TPD) (*64*). Our findings reveal a soluble antigen–gated process that drives clustering and degradation of a cell–surface receptor. To our knowledge, this is the first demonstration of ligand-dependent extracellular receptor degradation contributing to antibody efficacy in a clinical context. This finding may expand the scope of TPD and highlight how degradation-based mechanisms may enable new therapeutic opportunities in immuno-oncology and beyond.

Several limitations of our study should be noted. VEGF concentrations used in our assays were higher than those typically measured in tumors (*62*, *63*), though *in vivo* VEGF is continuously produced by tumor and stromal cells, making direct comparisons challenging. How these differences in concentration and exposure dynamics affect Ivonescimab activity remains an open question. In addition, *in vivo* evidence of Ivonescimab-driven PD-1 clustering and degradation is currently lacking, and studies to measure PD-1 levels in patient samples following Ivonescimab treatment would be highly informative. Finally, the effects of the antibody on VEGF signaling were not examined in our model. An interesting possibility is that VEGF may be co-degraded with PD-1, potentially effecting the tumor microenvironment through an additional mechanism that deserves further investigation.

In summary, Sphyder introduces a generalizable strategy for site-specific sensing and rewiring of tyrosine phosphorylation signaling. By creating modular phosphosite–binder pairs, Sphyder enables mechanistic dissection of signaling with spatiotemporal resolution, provides a framework for understanding responses to both established and emerging therapies, and establishes a platform for engineering synthetic immune signaling circuits. These advances lay the foundation for fundamental studies of immune regulation, engineering of biosensors to facilitate drug discovery, and the development of next-generation immunotherapies built on phosphosite-specific functional rewiring.

## Acknowledgements

We thank all members of the Zhou Lab for helpful discussions. We thank Aoxing Cheng for assistance with tumor cell line culture, plasmid construction, and protein purification, Jennifer Jiang for help with plasmid and LAT library construction, Kun Huang from the DFCI Molecular Imaging Core for guidance on confocal microscopy, and Milka Kostic and Marie Bao for their input on the manuscript.

## Author Contributions

Z.M. and X.Z. conceived the project and wrote the manuscript. Z.M. designed and performed all experiments. L.H. assisted with plasmid construction and contributed to figure and manuscript preparation. S.K.E. contributed to plasmid construction, library construction, and assay development. M.C.R. and R.G.M. provided CAR plasmids, LAT peptide, cell lines and assisted with human primary T cell studies. X.Z. supervised the study. All authors reviewed and approved the manuscript.

## Data and materials availability

The described constructs will be shared on reasonable request from academic researchers.

## Supplementary Figures

**Supplementary Figure S1.**
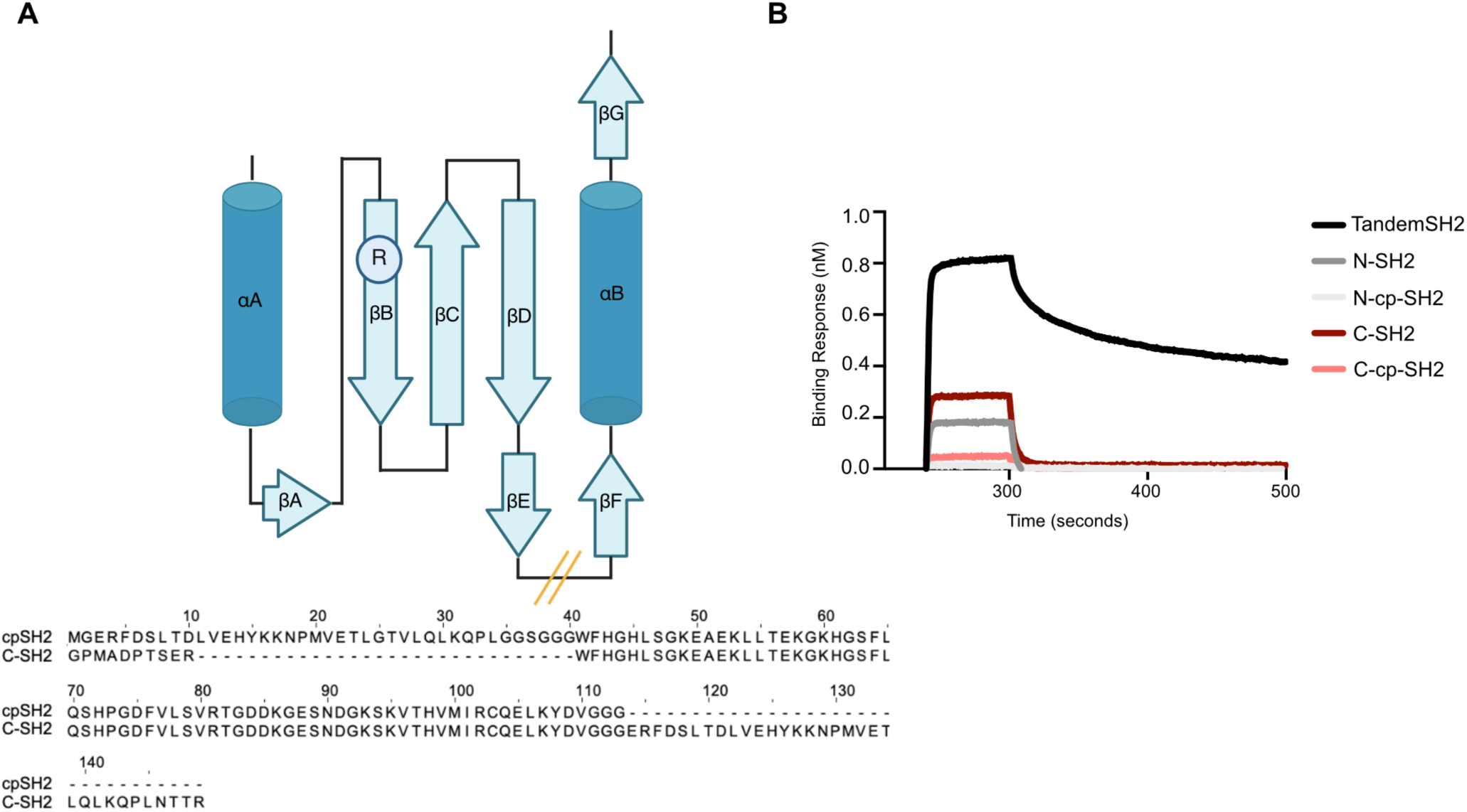
EF loop breakpoint selected as cut site for circular permutation of SHP2 SH2 domains. **(A)** Topological graph of SH2 domains, arginine on βB sheet is critical for its recognition of phosphotyrosine while others facilitate binding to phosphotyrosine contained linear motifs. Circular permutation of SH2 domain of SHP2 C-SH2 was performed by cutting the loop between βΕ and βF and connecting the C terminal with a GGSG linker to the original N terminal. **(B)** BLI curves showed dual SH2 domain from SHP2 binds to PD-1 pY248 containing peptide, while N and C terminal SH2 alone had a moderate affinity. Circular permutation of C-SH2 dampened the affinity further.

**Supplementary Figure S2.**
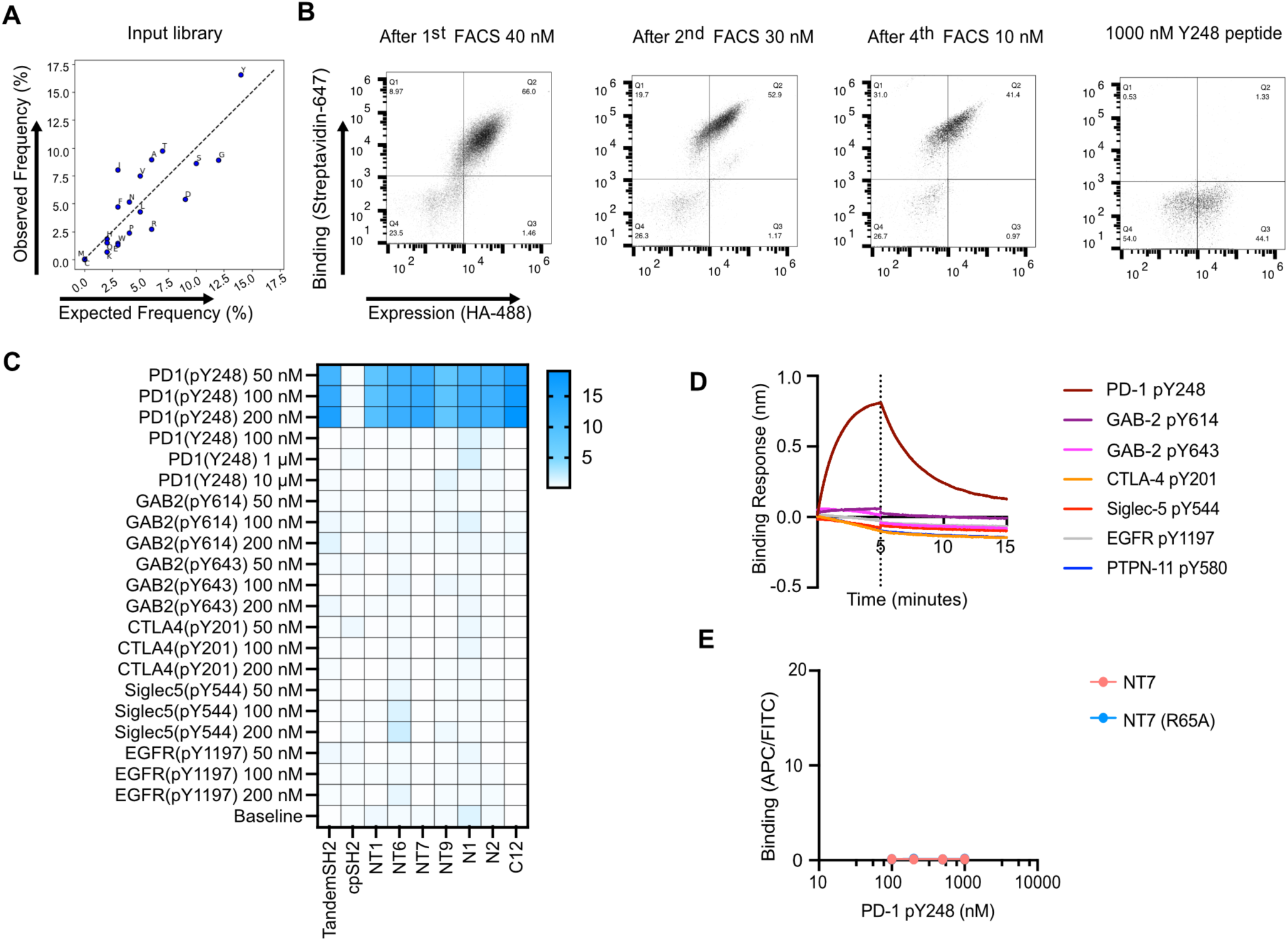
Engineering high-affinity Sphyder binders specific for PD-1 pY248. **(A)** Ampilcon sequencing was performed in the starting input library, the observed amino acid composition of the CDRs was comparable with expected frequencies, and throughout selection the composition remained similar to expectations.**(B)** Yeast surface display screening of a library encoding SHP2 C-cpSH2 fused to a Nb for binding to biotinylated PD-1 pY248 peptide. Flow cytometry plots show progressive enrichment of high-binding clones across four rounds of FACS using decreasing peptide concentrations (from 40 nM to 10 nM). A final negative selection with 1000 nM unphosphorylated Y248 peptide removes clones recognizing non-phosphorylated sequences. **(C)** Heatmap showing on-yeast specificity profiling of top Sphyder binders against a panel of phosphotyrosine peptides from diverse immune signaling proteins. Binders show strong and selective binding to PD-1 pY248 peptide while exhibiting minimal cross-reactivity to other phosphopeptides. **(D)** BLI curves showing binding kinetics of recombinant NT7 binder against phosphorylated PD-1 pY248 peptide (red trace) and various unrelated phosphotyrosine peptides (colored traces). NT7 binds PD-1 pY248 with nanomolar affinity and exhibits negligible binding to non-cognate phosphopeptides.

**Supplementary Figure S2.**
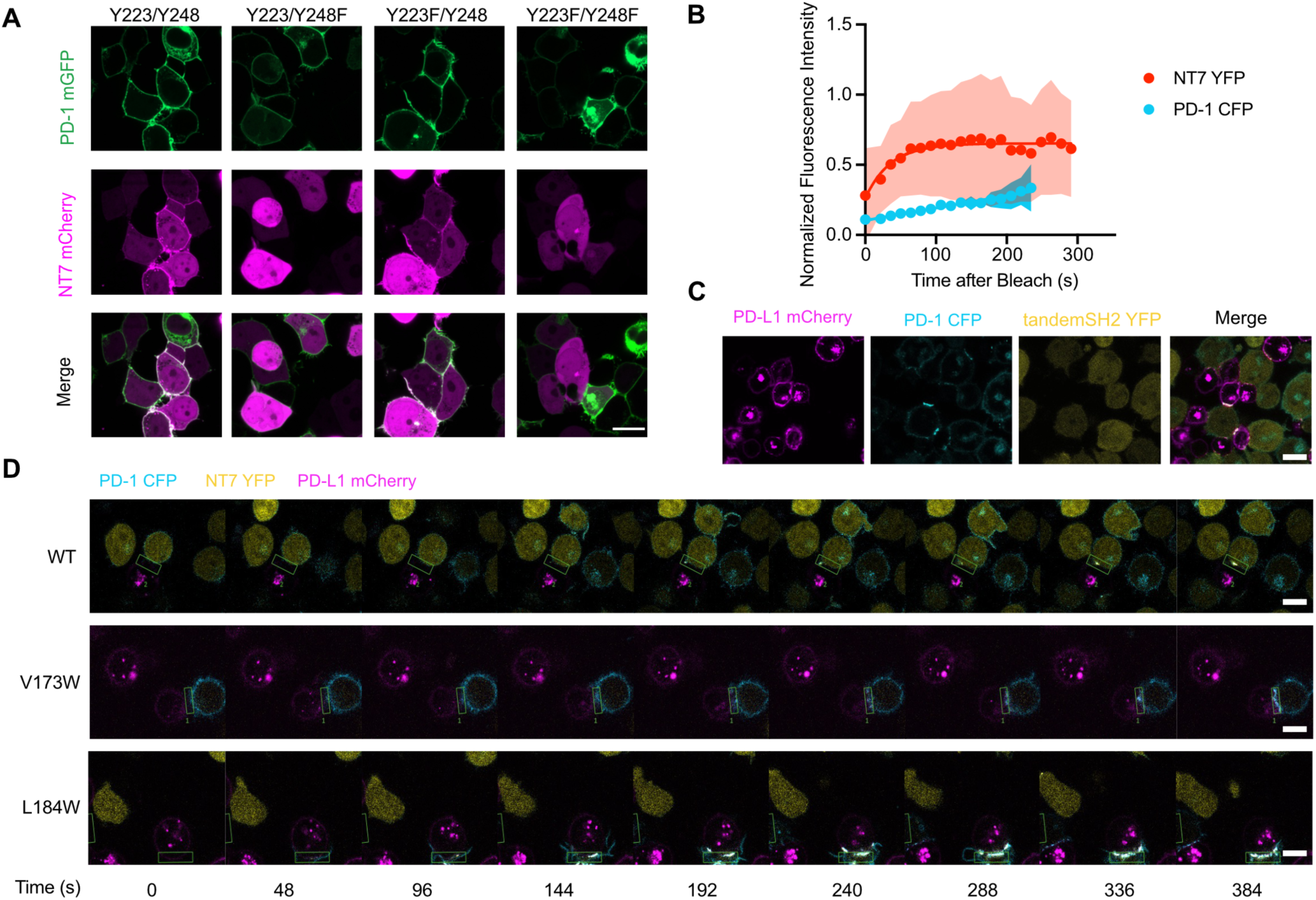
Dimerization-dependent regulation of PD-1 phosphorylation visualized with Sphyder sensors. **(A)** Confocal images of HEK293 cells co-expressing NT7-mCherry and various PD-1-GFP constructs containing wild-type or mutant intracellular tyrosine residues (Y223/Y248, Y223F/Y248, Y223/Y248F, Y223F/Y248F). NT7-mCherry localizes to the plasma membrane only in cells expressing PD-1 constructs with Y248 but not F248, confirming pY248 binding. Scale bars, 10 μm. **(B)** Fluorescence recovery after photobleaching (FRAP) analysis of NT7-YFP and PD-1-CFP signals. NT7-YFP recovers rapidly after bleaching, indicating dynamic binding to phosphorylated PD-1, while PD-1-CFP shows slower recovery. Data are normalized to pre-bleach fluorescence intensity. **(C)** Confocal images of PD-L1-mCherry+ Raji cells in co-culture with PD-1-CFP+, tandem SH2-YFP+ Jurkat T cells. Unlike NT7, the tandem SH2 construct shows minimal co-localization with PD-1 at the immune synapse. **(D)** Time-lapse imaging of Jurkat T cells expressing wild-type, V173W, or L184W PD-1 mutants, showing dynamic localization of PD-1-CFP, pY248 binder-YFP, and PD-L1-mCherry over time during immune synapse formation. The L184W mutant exhibits higher Sphyder binder recruitment, consistent with increased phosphorylation due to enhanced PD-1 dimerization. The V173W mutant displays reduced recruitment, indicating impaired phosphorylation dynamics. Time stamps shown in seconds.

**Supplementary Figure S4.**
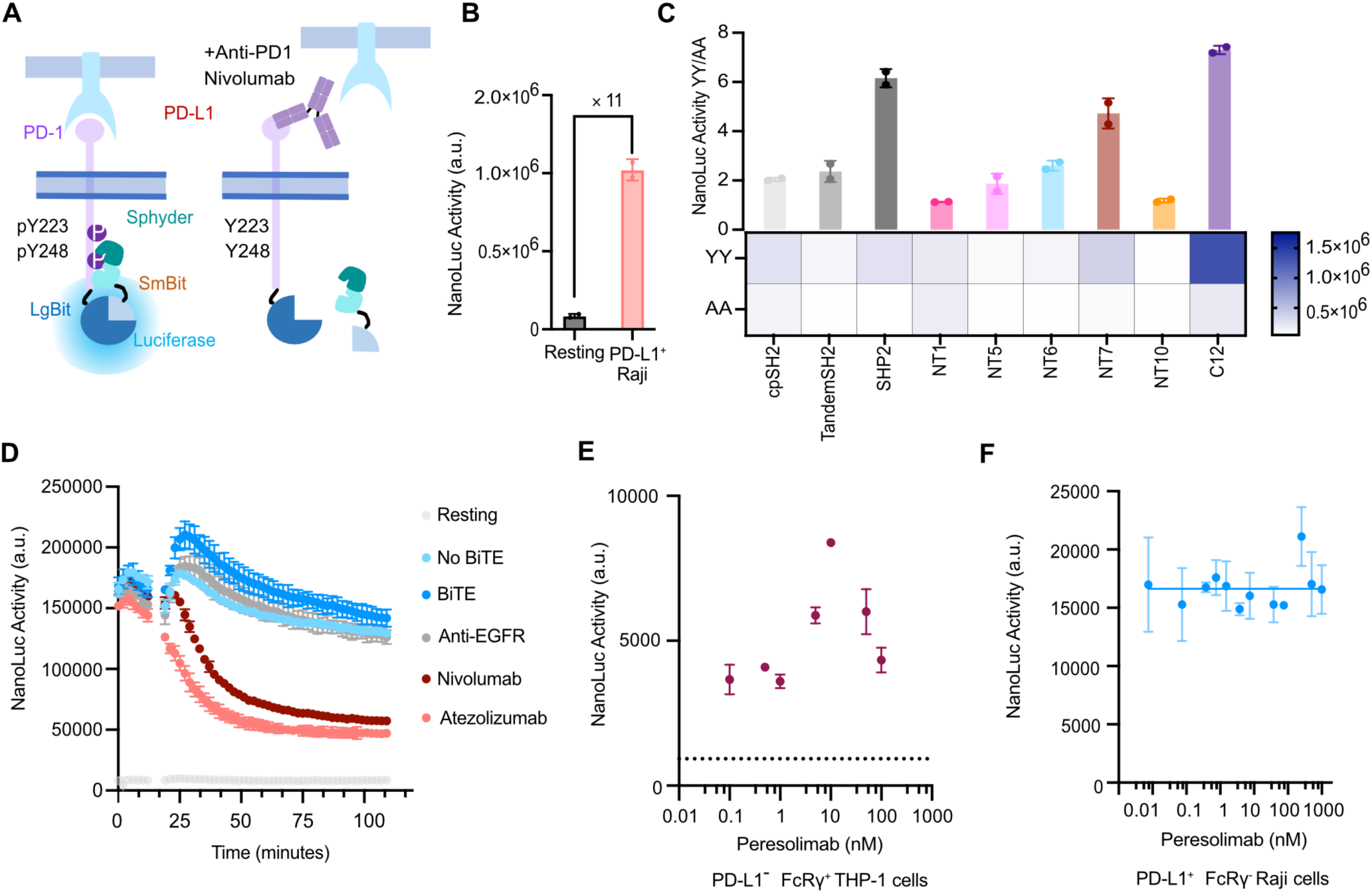
Quantitative live-cell luciferase assays for measuring PD-1 phosphorylation and response to immune checkpoint modulators. **(A)** Schematic of the split NanoLuc luciferase assay for monitoring PD-1 phosphorylation in living cells. In resting cells (left), PD-1-LgBiT and NT7-SmBiT remain spatially separated. Upon PD-1 phosphorylation, NT7-SmBiT is recruited to phosphorylated PD-1, enabling reconstitution of the NanoLuc enzyme and generating a luminescence signal. Treatment with anti-PD-1 ICIs, such as nivolumab (right), prevents PD-1 phosphorylation and reduces NanoLuc signal. **(B)** Bar graph showing NanoLuc luciferase activity of PD-1 Split-NanoLuc cells in resting state and co-culture with PD-L1 expressing Raji cells. **(C)** Bar graph showing NanoLuc luciferase activity in PD-1-LgBiT⁺ NT7-SmBiT⁺ Jurkat cells expressing either wild-type PD-1 (Y223/Y248; “YY”) or tyrosine-mutated PD-1 (Y223A/Y248A; “AA”). Several engineered Sphyder binders, including NT7 and C12, show high selectivity for phosphorylated PD-1, confirming phosphorylation-dependent binding. Data represent mean ± s.d. from 2 biological replicates. **(D)** Time-course luminescence measurements in PD-1 Split-NanoLuc Jurkat cells co-cultured with PD-L1⁺ Raji cells under different conditions. Addition of a CD3/CD19 BiTE induces strong PD-1 phosphorylation and luciferase activation (blue trace). This signal is rapidly reduced upon treatment with anti-PD-1 (nivolumab) or anti-PD-L1 (atezolizumab) ICIs, indicating effective inhibition of PD-1 signaling. Anti-EGFR antibody is included as an irrelevant control. **(E)** Dose-response curve for peresolimab, a clinical-stage PD-1 agonist antibody, tested in PD-1 Split-NanoLuc Jurkat cells co-cultured with PD-L1⁺ THP-1 cells. Peresolimab induces a dose-dependent increase in luciferase activity at concentrations between 0.1 and 10 nM, with diminished effects at higher doses. **(F)** Dose-response curve for Peresolimab in FcRγ non expressing PD-L1 expressing Raji cells, no agnostic effect could be observed.

**Supplementary Figure S5.**
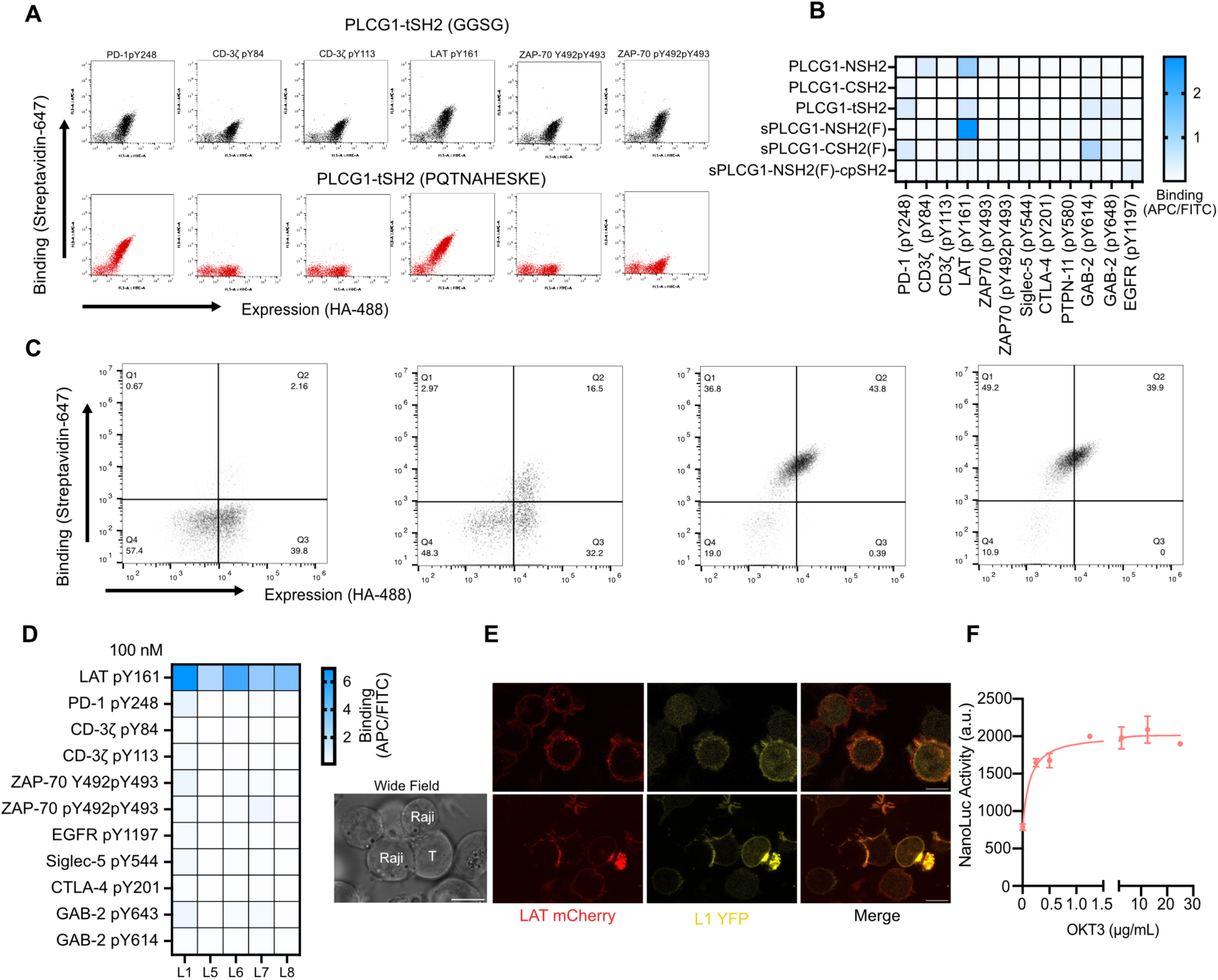
Characterization of Sphyder binders for LAT pY161. **(A)** Flow cytometry analysis of yeast-displayed PLCG1 tandem SH2 binding to various phosphopeptides, including PD-1 pY248, CD3ζ ITAM pY83 and pY111, LAT pY191, and ZAP70 pY492/pY493. Upper panels show binding of the PLCγ1-SH2 domains with a flexible GGSG linker; lower panels show binding of the PLCγ1-SH2 domains with endogenous linker (PQTNAHESKE). **(B)** Heatmap representation of phosphopeptide binding specificity for different PLCG1 SH2 constructs. Binding was measured as APC/FITC ratio across phosphopeptide targets, showing selective enrichment for LAT pY161 by superbinder constructs. **(C)** Representative flow cytometry dot plots showing expression levels (HA-488) on the x-axis and phosphopeptide binding (streptavidin-647) on the y-axis for yeast-displayed LAT binder candidates. Data demonstrate enrichment of high-binding clones. **(D)** Heatmap summarizing specificity of enriched LAT binders (L1–L8) against a panel of phosphopeptides. Signal intensities indicate selective binding to LAT pY161, with negligible binding to unrelated phosphopeptides such as PD-1 pY248, CD3ζ ITAMs, EGFR, CTLA4, and GAB2. **(E)** Confocal microscopy images of Jurkat T cells co-cultured with Raji cells, showing LAT-mCherry localization (red), co-localization with LAT pY161 Sphyder binder fused to YFP (yellow), and merged channels. Sphyder-LAT binder specifically localizes to LAT at the immune synapse, demonstrating intracellular recognition of phosphorylated LAT pY161. Scale bar, 10 μm. **(F)** A split luciferase assay of LAT activation, where LAT was fused with LgBit and co-expressed with Sphyder-LB1-SmBit. Reporter expressed Jurkat cells were cultured in an anti-CD3 (OKT3 clone) coated plated for 15 minutes, luminescence was measured with Vivazine. NanoLuc activity was dose responsive to anti-CD3, suggesting LAT activation. Data represent mean ± s.d. from 2 replicates.

**Supplementary Figure S6.**
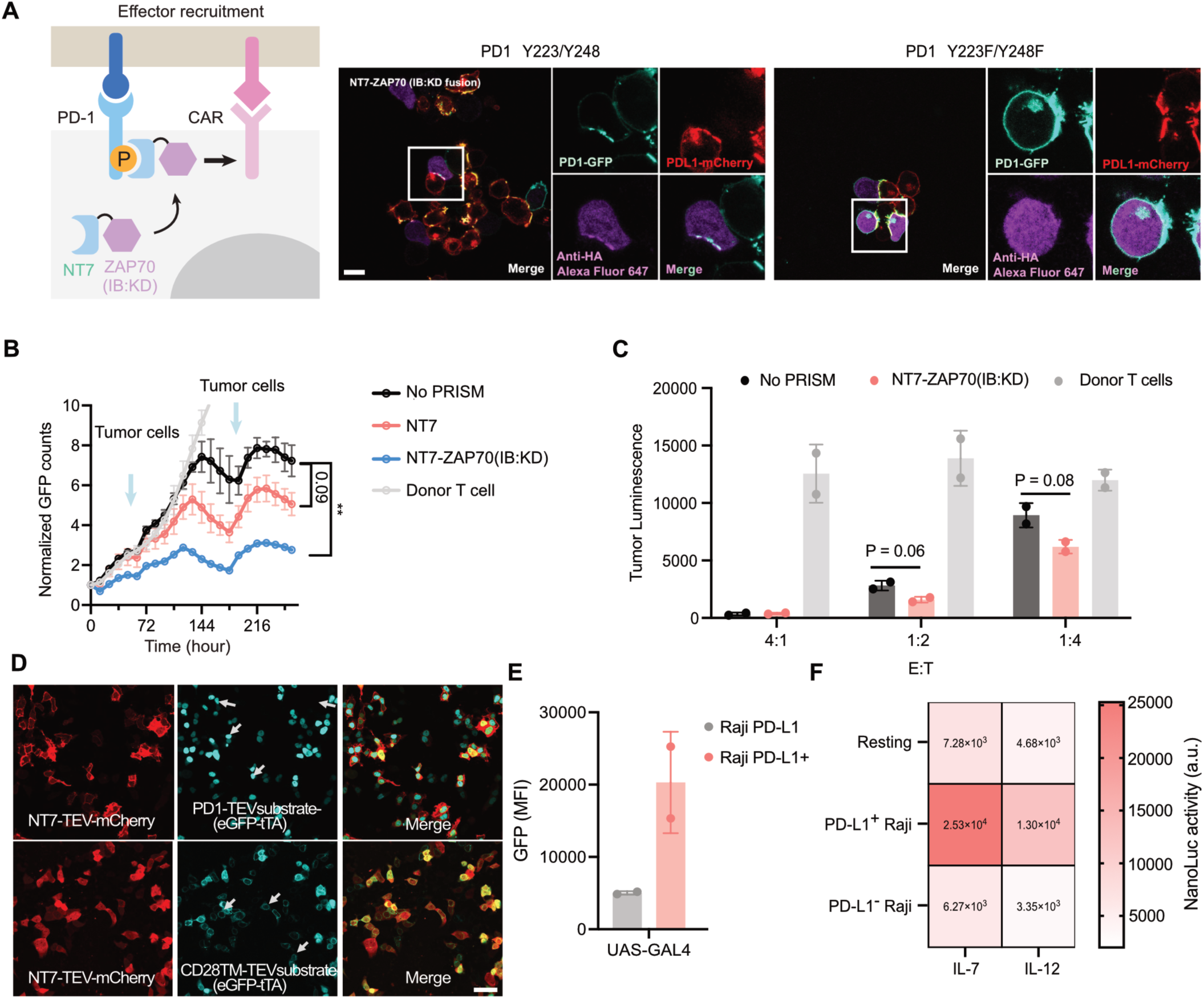
Synthetic rewiring of PD-1 signaling. **(A)** Engineered PD-1 pathway using Sphyder binders. Phosphorylated PD-1 recruits synthetic modules that convert inhibitory signaling into immune activation via kinase recruitment, NT7 fused to the interdomain B and kinase domain of ZAP70 (IB:KD) redirects kinase activity to phosphorylated PD-1. Confocal imaging of NT7-ZAP70(IB:KD) recruitment to phosphorylated PD-1 in Jurkat T cells. Wild-type PD-1 recruits the fusion protein to the membrane (up), while the phosphorylation-deficient mutant does not (down). Scale bars, 10 μm. **(B)** Quantification of GFP-expressing SY5Y tumor cells in a repeat-challenge assay using primary human GD2 HA-28z CAR-T cells expressing various Sphyder constructs. Two-way ANOVA, ** P=0.0018. **(C)** PD-L1^+^ SY5Y cells expressing firefly luciferase were co-cultured with Sphyder-NT7-ZAP70(IB:KD) armed CAR-T cells in different Effector:Tumor cells ratio, luminescence signal was then measured after 72 hours of co-culture. **(D)** In HEK293T cells, PD1-TEVsubstrate-eGFP-tTA, CD28TM-TEV-substrate-eGFP-tTA were co-expressed with Sphyder-NT7-TEV-mCherry. Membrane localization and nuclear translocation of eGFP-tTA could be observed in PD1 expressing cells but not CD28TM anchored eGFP-tTA cells. Scale bars: 50 μm. **(E)** PD-1 circuit could use other transcription factors such as GAL4-VP64 which also showed induction of UAS::GFP expression when co-cultured with PD-L1^+^ Raji cells. **(F)** Jurkat cells armed with PD-1 tTA circuit driven IL-2, IL-7 and IL-12. Co-culture with PD-L1 expressing Raji cells showed robust induction of all three cytokines, while co-culture with PD-L1 non expressing Raji cells, minimal cytokine induction was observed. BiTE could induce the cytokine expression more substantially in co-culture with Raji PD-L1^+^ compared with PD-L1 non expressing cells.

**Supplementary Figure S7.**
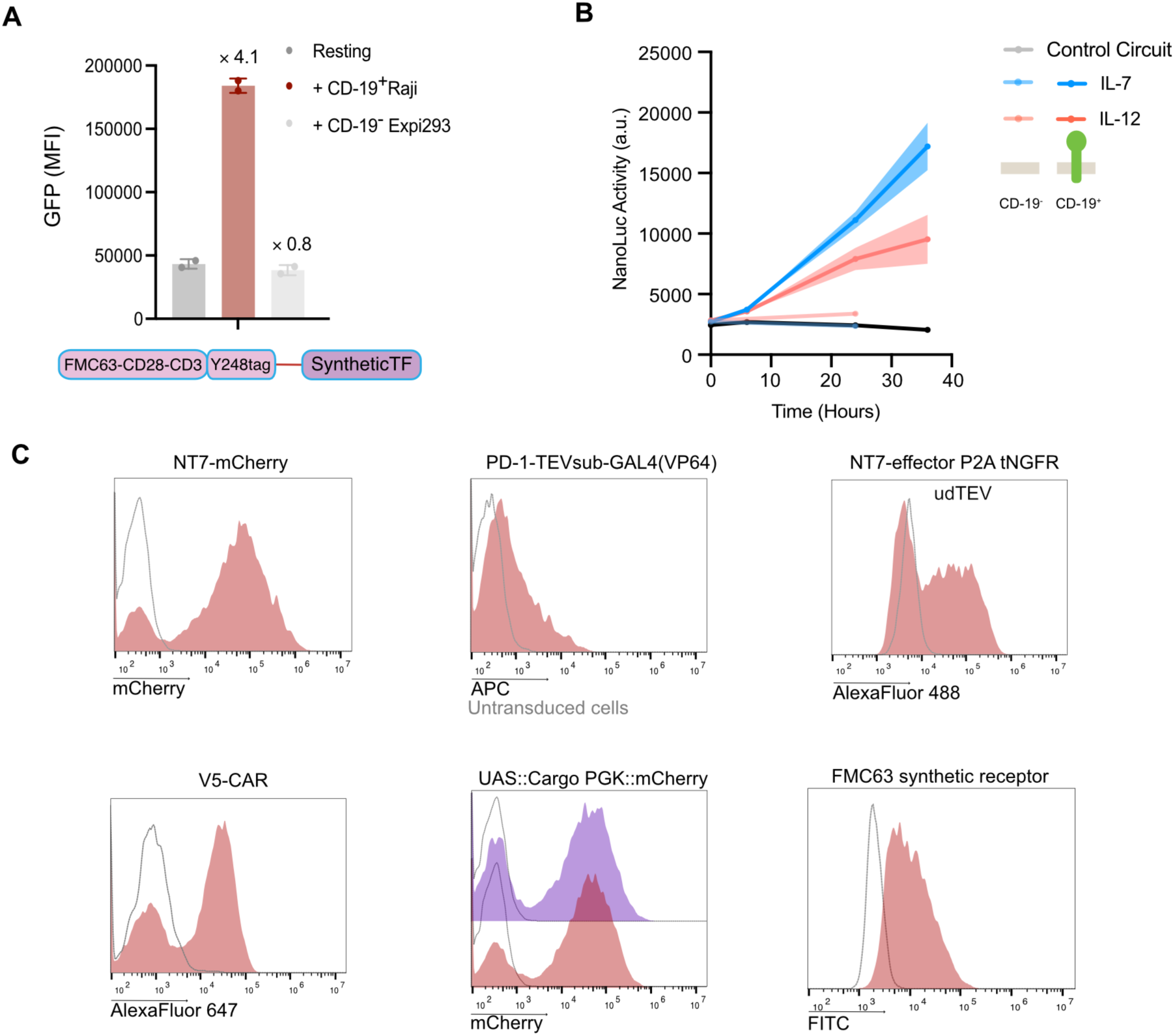
Modular reprogramming of PD-1 signaling with PRSIM-NT7. **(A)** TEV substrate fused TF was linked to a PD-1 Y248 site fused CD19-28 CAR. Expression of this fusion CAR induced GFP expression in Jurkat cells when co-cultured with CD19^+^ Raji cells. Data represent mean ± s.d. from 2 replicates. **(B)** A CD-19 targeting Sphyder circuit showed time dependent and antigen dependent secretion of cytokine cargos measured with split NanoLuc where cytokine cargos were tagged with HiBit. CD-19 targeting Sphyder circuit expressing primary T cells were co-cultured with CD-19 expressing Nalm6 cells, supernatant was harvested according to the time points for analysis. **(C)** Representative flow cytometry analysis of expression of Sphyder-NT7 and Sphyder-NT7 fusion protein, PD-1 TF fusion, CAR, synthetic TF response elements and FMC63 ectodomain replacement chimeric receptor in human primary T cells.

**Supplementary Figure S8.**
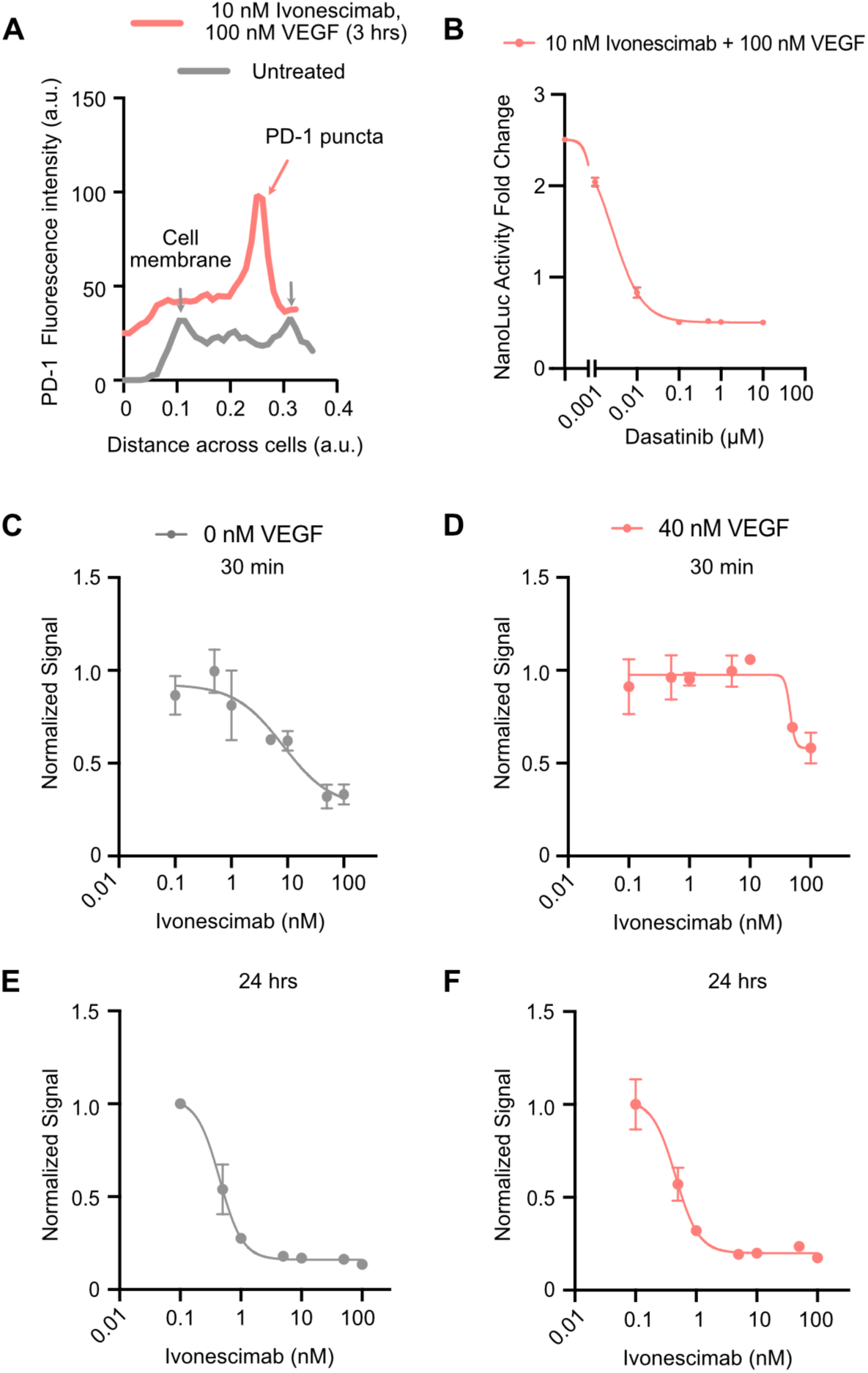
Ivonesicmab drives clustering mediated phosphorylation, endocytosis and degradation of PD-1 on the cell surface. **(A)** Representative quantification of PD-1 puncta fluorescence corresponding to Fig. 5G. In untreated cells, PD-1 localized primarily to the plasma membrane (gray arrows). In cells treated with 10 nM Ivonescimab and 100 nM VEGF for 3 h, prominent PD-1 clusters formed (red arrow). For consistency, puncta were aligned to the right side in the quantification plots when asymmetrically distributed. **(B)** Normalized luciferase activity in Split-NanoLuc Jurkat cells treated with increasing concentrations of LCK-inhibitor Dasatinib. Cells were treated for 30 min with 10 nM Ivonescimab and 100 nM VEGF. **(C-F)** ormalized NanoLuc luciferase activity in Split-NanoLuc Jurkat cells co-cultured with PD-L1⁺ Raji cells plus 1 nM BiTE, treated with increasing doses of Ivonescimab for 30 min (C–D) or 24 h (E–F). VEGF concentrations: 0 nM (C, E), 40 nM (D), and 50 nM (F). At 24 h, Ivonescimab acted as a potent antagonist with a sub-nM IC_50_. At 30 min, Ivonescimab showed reduced potency, particularly in the presence of VEGF. Data represent mean ± standard deviation from n = 2 replicates.

**Supplementary Table S1.**
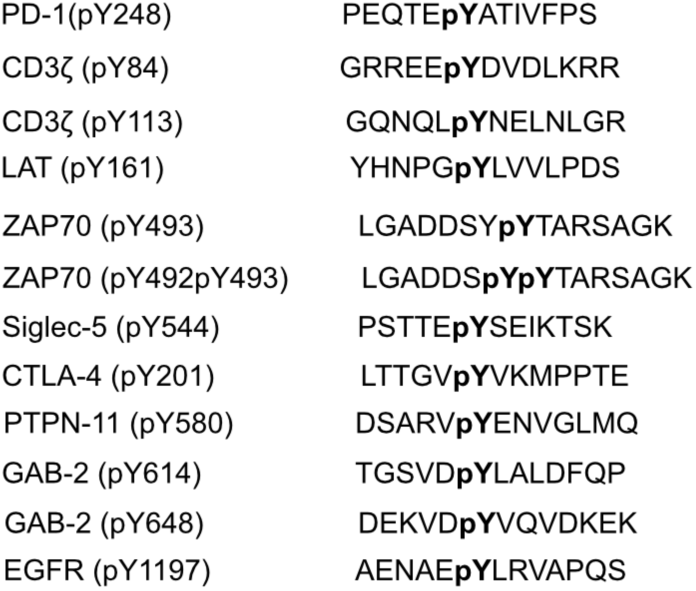
Sequences of phosphotyrosine containing peptides.

